# A systematic literature review of forecasting and predictive models of harmful algal blooms in flowing waters

**DOI:** 10.1101/2025.09.29.679270

**Authors:** Jennifer C. Murphy, Rebecca M. Gorney, Lisa A. Lucas, Jacob A. Zwart, Jennfier L. Graham

**Affiliations:** U.S. Geological Survey, Central Midwest Water Science Center; U.S. Geological Survey, New York Water Science Center; U.S. Geological Survey, Water Mission Area

**Keywords:** Cyanobacteria, rivers, harmful algal blooms, modeling, data-driven, process-based

## Abstract

Occurrences of harmful algal blooms (HABs) in rivers challenge the belief that rivers are not susceptible to HABs because of their short residence times and fluctuating hydrology. Here we present a systematic literature review of predictive and forecasting models for HABs in flowing waters, including rivers, flowing in-stream reservoirs (e.g., run-of-river reservoirs and lock-and-dam systems) and tidal or estuarine systems with riverine processes. The review aimed to understand current and historical modeling approaches for predicting and forecasting river HABs, without restricting to specific taxa, such as cyanobacteria, or modeling endpoints. The review included 162 articles published over nearly 50 years, covering more than 80 rivers worldwide. Eutrophic, non-wadable rivers with in-stream obstruction were commonly modeled, though diverse environmental characteristics were reported. Most articles used algal biomass or chlorophyll as modeling endpoints, with a quarter using novel or unique endpoints. Algal toxins motivated model development in 23% of the articles, however just 5% used algal toxins as an endpoint. Only 6% of the articles modeled benthic HABs; the rest focused on pelagic HABs. There was no standard model used for modeling river HABs. Process-based models were more common (59%) than data-driven approaches (37%), with model formulations ranging from simple to complex, which contrasts with a lake-focused literature review of HAB models that found data-driven models were more common. Models in river settings shared similar input variables as those previously identified for lakes, such as water temperature, nutrients, and light availability. However, streamflow and other transport metrics took prominence in river models compared to lake models. Algal cell physiology (such as growth, predation, and motility) was routinely included as input data or as mathematical formulations in process-based models and these processes were frequently identified as an important predictor by the articles’ authors. Conversely, data-driven models rarely included these processes, instead using predictors related to environmental conditions, such as nutrients, water quality, water temperature, and streamflow. These important proxy predictors have apparent success with modeling overall algal biomass (irrespective of taxa) whereas other factors, such as those related to algal physiology and other biological processes, are likely responsible for more subtle shifts in community composition. These differences highlight the influence of data availability, especially for processes that are difficult, time-consuming, or expensive to measure, on model development and model outcomes, raising questions about the selection of modeling inputs and endpoints. Challenges to advancing river HAB modeling include the lack of site-specific model inputs representing key processes (e.g., photosynthetic parameters and predation rates), overlooked riverine environments like the benthos and side/back-channel areas, lack of information on environmental settings, and poorly reported model performance metrics. This review emphasizes opportunities for advancing river HAB modeling by learning from well-honed estuarine models, supporting current forecasting and operationalization efforts, and developing common datasets for river HAB model development and evaluation.

## 1. Introduction

Harmful algal blooms (HABs) are a global phenomenon that occur in diverse aquatic ecosystems spanning the freshwater to marine continuum (Glibert, 2017; Howard et al., 2023; Peacock et al., 2018; Stauffer et al., 2019). The potential harms associated with blooms include excess algal biomass, degraded water quality, excessive oxygen demand, disrupted aquatic food webs, and production of secondary metabolites, such as taste and odor causing compounds and toxins, all of which may have far-reaching ecologic, economic, and public health consequences (Brooks et al., 2016; Chorus and Welker, 2021; Huisman and Weissing, 1994). The organisms responsible for HABs and the environmental conditions that foster their development arise from a complex interplay of physical, chemical, and biological processes that occur across various spatial and temporal scales (Burford et al., 2020; Glibert, 2017; Griffith and Gobler, 2020; Zhou et al., 2020).

In freshwaters, prokaryotic cyanobacteria are the primary organisms that cause HABs and are the only freshwater taxa known to produce toxins that can adversely affect human health. However, other freshwater eukaryotic algae (e.g. diatoms and green algae) can cause HABs and produce toxins (e.g., chrysophytes and euglenophytes) that affect aquatic organisms, particularly fish (Gorney et al., 2023; Patiño et al., 2023). Key environmental factors that influence algal community composition and bloom development include water temperature, which affects algal physiology and growth; light, essential for photosynthesis; and nutrients, crucial for cellular function (Chorus and Welker, 2021; Patiño et al., 2023). Physical processes are also important, and in lotic environments hydrodynamic processes moderate the effects of water temperature, light, and nutrients on algal growth (Cha et al., 2017; Chételat et al., 2006; Graham et al., 2020; Reynolds and Descy, 1996; Van Nieuwenhuyse and Jones, 1996). Any specific location along a river is inherently connected with upstream physical, chemical, and biological processes (Glibert, 2017; Junk et al., 1989; Thorp et al., 2006; Vannote and Sweeney, 1980; Walker et al., 2006). Ephemeral connectivity to backchannel areas (Giblin and Gerrish, 2020; Giblin et al., 2022), reservoir releases and flow control (Graham et al., 2012; Otten et al., 2015; Williamson et al., 2018), and downstream transport from upstream zones of benthic or pelagic productivity (Schmadel et al., 2024; Wood et al., 2020) may all contribute to HABs in lotic environments. Consequently, the negative effects of freshwater HABs can extend hundreds of miles downstream from upstream source areas and eventually impair estuarine and coastal environments (Miller et al., 2010; Peacock et al., 2018; Preece et al., 2017).

Models serve as important tools to enhance our understanding of HABs and aid in forecasting and management decisions. HAB modeling is performed across various spatiotemporal scales, from examining the physiological responses of specific taxa in laboratory conditions to analyzing global drivers and simulating long-term changes in occurrence (Burford et al., 2020; Lucas and Deleersnijder, 2020; Reynolds, 1998; Stauffer et al., 2019). Modeling efforts inform public health protection, mitigation strategies, and resource management. Early indicators and near-term event forecasts allow proactive responses to potentially hazardous conditions, thereby safeguarding animal and human health and reducing economic losses (Petchey et al., 2015). Predictive modeling can help assess outcomes of future scenarios, including management strategies under changing environmental conditions, ideally supporting informed decision making. Therefore, investment in the development of robust forecasting and predictive models has been recognized as a critical need in addressing the challenges posed by HABs in freshwaters (Burford et al., 2020; Rousso et al., 2020; U.S. National Office for Harmful Algal Blooms, 2024).

Rousso et al. (2020) systematically reviewed literature on forecasting and predictive models for harmful cyanobacterial blooms (CyanoHABs) in freshwater lakes and reservoirs. Their review revealed that most lake models were site- and species-specific, primarily focused on nutrient-enriched systems, and had inconsistent predictor variables across models; nonetheless, water temperature, phosphorus, and nitrogen were consistently identified as important predictor variables across various model types. These findings highlight the complexity of cyanobacterial community dynamics and bloom formation. The development of CyanoHAB models has paralleled advancements in computational capabilities and in monitoring technologies, such as machine learning, high-frequency sensors and remote sensing. Rousso et al. (2020) noted that challenges in comparing the performance of different models arise due to variability and inconsistent reporting of location and frequency of sampling, analytical measurement procedures, and monitoring duration, alongside a lack of consistent model performance metrics. A key conclusion of the review was the necessity for establishing a CyanoHAB modeling database; the compilation of lake studies reviewed by Rousso et al. (2020) serves as a foundational dataset for such an initiative. In contrast, Xia et al. (2019) offer a qualitative, albeit non-systematic, literature review of algal blooms in large rivers aiming to define river blooms, describe their negative effects, and identify likely key drivers. Although, they present a useful conceptual framework for understanding blooms in these systems, the study does not provide a quantitative comparison of the literature nor an evaluation of different modeling approaches and processes. Given the groundwork laid by Rousso et al. (2020) and Xia et al. (2019), a systematic literature review of HAB models in lotic settings would provide a point of comparison to similar modeling in lake settings and potentially improve forecasting capabilities and provide insights into future HAB conditions across freshwater systems.

We adapted the methods outlined by Rousso et al. (2020) to systematically review the current literature for HAB forecasting and predictive models for riverine environments for a more comprehensive view of HAB modeling in freshwaters. In alignment with their definitions, we distinguish between forecasting and predictive models versus models used for validation/qualitative explorations of observed data. Forecasting and predictive models provide future estimates focused on informing short-term operational strategies, long-term projections used for scenario analysis or estimates between observations temporally or spatially. We included models used for sensitivity analyses, hindcasting or nowcasting, which were not included in Rousso et al. (2020). Our review compiled articles with models that estimated HAB-related variables at locations and (or) times not represented in calibration or training datasets. As such, like Rousso et al. (2020), we excluded models that solely analyzed and interpreted empirical data, despite the valuable insights they may provide. Although Rousso et al. (2020) focused exclusively on cyanobacteria, our review encompassed all freshwater taxa associated with potentially harmful blooms. We organized our findings into four thematic areas: environmental setting, modeling data, model types, and model application. Within each theme, we present both quantitative and qualitative summaries of the literature and highlight advancements and insights that were beneficial to researchers. Finally, we identify challenges facing the river HAB modeling community and suggest key opportunities for advancing river HAB modeling, particularly in the context of HAB management and mitigation.

## 2. Methods

The systematic literature review was completed in three phases: (1) a literature search of major scientific databases, (2) a three-step screening process to identify a final set of articles, and (3) extraction of information from each article during a critical review (Figure 1). We largely followed the steps laid out by Rousso et al. (2020) and described by Pickering and Byrne (2014). The benefit of a systematic literature review is that well-defined search queries and inclusion/exclusion criteria make the review reproducible and less dependent on the area of expertise of the researchers completing the literature review.

**Figure 1.**
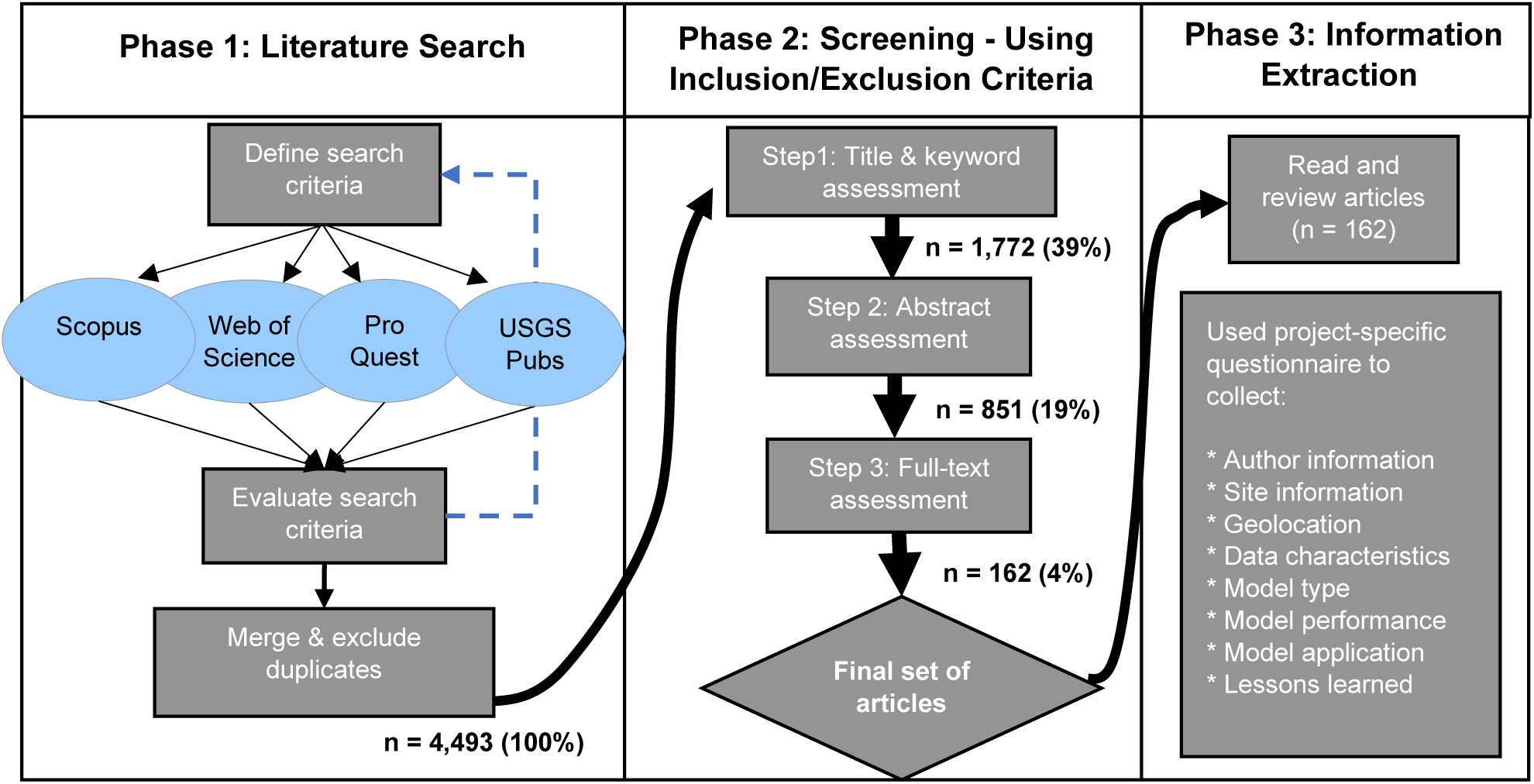
Workflow for the systematic literature review, including the number (n) and percentage (%) of articles retained after each step. We searched Scopus, Web of Science, ProQuest, and U.S. Geological Survey (USGS) publications (USGS Pubs) in the Phase 1 literature search. The blue dashed line reflects iterative refinement of Phase 1 search criteria to ensure validation papers were captured. This workflow was adapted from that of Rousso et al. (2020).

### 2.1 Phase 1 – Search queries and literature sources

During Phase 1 of the literature review, we developed a set of queries to use as search criteria in databases of scientific, peer reviewed publications. The search criteria included four categories of information (Table 1) used to target publications focused on freshwater algae under bloom conditions, in river settings, that developed or applied predictive or forecasting models. The four queries were used together, connected with the “and” (i.e., “&”) Boolean operator. Queries were refined over several iterations. We used a set of 24 validation papers (i.e., papers we knew contained river HAB models; Table SM-1) which we were able to retrieve using the final search criteria (Table 1).

**Table 1.**
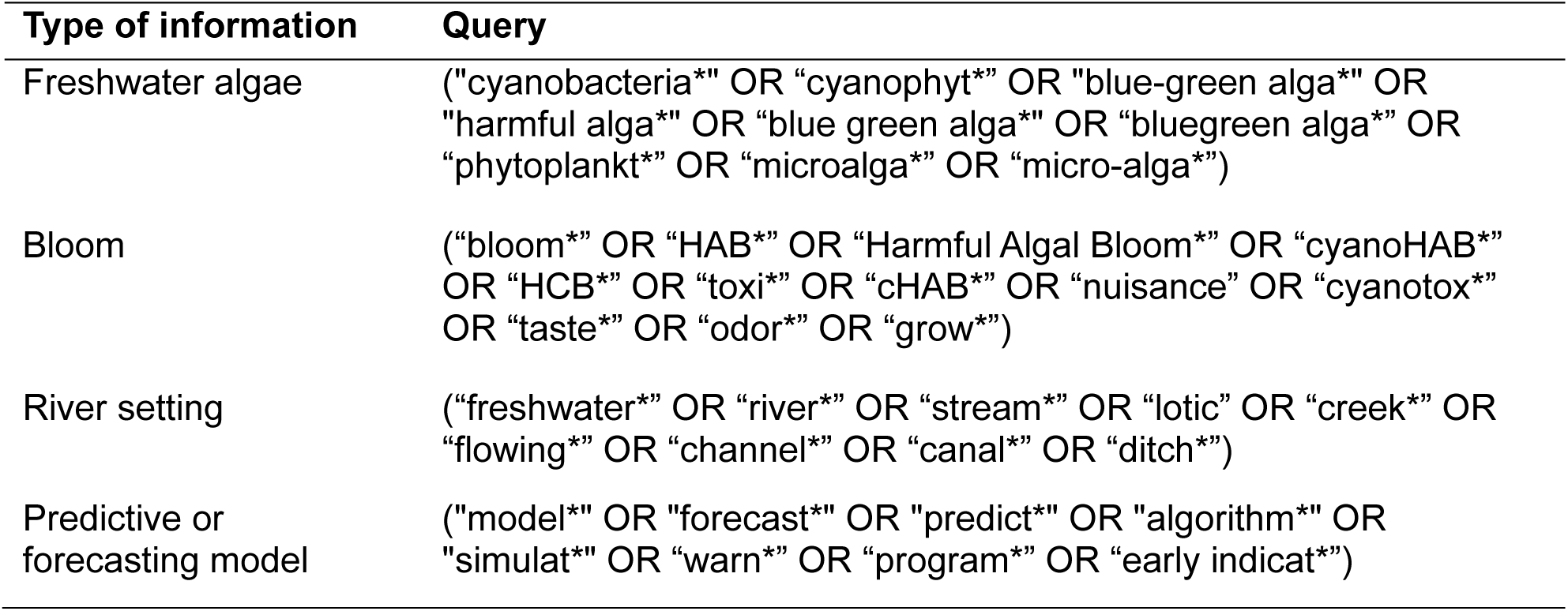
Search criteria applied in Phase 1 of systematic literature review. All queries were used together with an “and” (&) operator. * is a wildcard that represents one or more additional characters.

In the spring of 2023, we applied the search criteria to titles, keywords, and abstracts (when available) in four scientific databases: Scopus, Web of Science, ProQuest, and the U.S. Geological Survey (USGS) Publications Warehouse. We merged results from all four database searches and removed duplicate entries. Phase 1 yielded the titles, keywords, and abstracts for 4,493 articles meeting our search criteria (Table 1; Figure 1).

### 2.2 Phase 2 – Inclusion/exclusion criteria and screening steps

Phase 2 involved application of inclusion/exclusion criteria (below) in a three-step screening process to identify the final set of papers to be critically reviewed in Phase 3 (Figure 1). First, we assessed the title and keywords of each article, then the abstract, and finally the entire article to determine whether an article satisfied the inclusion/exclusion criteria. If it became clear, after screening the title and keywords or abstract, that a paper did not satisfy the criteria, the succeeding screening step(s) were not necessary. A paper needed to meet all three requirements of the inclusion/exclusion criteria and clear all exclusions to be included in the final set. The full inclusion/exclusion criteria used during Phase 2 are provided in Table SM-2 in Supporting Materials-1. In brief, the inclusion and exclusion criteria stipulated:

1. The research must have been conducted in freshwater, flowing environments such as rivers, streams, creeks, canals, channels, and ditches, whether natural, constructed, modified, or managed. Run-of-river reservoirs, lock and dam pools, and rivers in estuarine settings were also included if the corresponding model contained clearly riverine processes (e.g.,

advective flow). Models of both real and idealized settings (i.e., observational data were not used and instead concepts of the environment were simplified or “idealized” for model development) were included.

1. Models must have been developed or implemented for prediction (i.e., between observations over time or across space), hindcasting, nowcasting, forecasting, or scenario or sensitivity analyses (e.g., related to environmental change or management decisions).
2. Modeling endpoints must have been measures of algal composition, abundance, biomass, presence, or toxicity (e.g., community composition, cell counts, biovolume, toxin concentrations) or related proxies (e.g., chlorophyll, phycocyanin). This included models that predicted primary production and algal cellular nutrient content. Novel proxies were also considered (e.g., metabolism metrics or oxygen dynamics) as long as the stated purpose was for algal bloom prediction or forecasting. Models providing categorical output or estimates of bloom risk or probability were also included.

Using the inclusion/exclusion criteria (Table SM-2), we retained 1,772 articles (39% of the initial 4,493 articles) after the title and keyword screen, 851 (19%) after the abstract screen, and finally 162 articles after the full text screen, which represented just 3.6% of all the articles returned from Phase 1 (Figure 1). We ensured that the final set included only peer reviewed, full-text, peer-reviewed articles written in English. We did not include conference proceedings, book chapters, or proposals.

### 2.3 Phase 3 – Information extraction

After completion of Phase 2, each of the 162 articles was critically reviewed. To assist in the extraction of information from each article, we developed a fillable online form using ArcGIS Survey123 (Esri, 2025) with a standard set of questions. A dataset containing the questions used in the form and the information extracted from each article is available in Gorney et al. (2025). We exported results from our critical review to a comma separated values (CSV) file and prepared and analyzed the data using the R statistical software (R Core Team, 2025).

Information on the following topics was retrieved for each article:

1. **Publication information**: Year of publication, affiliation of author, publication outlet
2. **Location information**: Name of system, geographic location, field setting, qualitative size of river, pelagic or benthic focus, and noteworthy environmental conditions described by the authors (e.g., eutrophic conditions, managed flows, point sources, etc.)
3. **Model information**: Model type(s) (data-driven, process-based, or other), how many models were developed or used, skill metric(s), and qualitative evaluation of skill

a. If data-driven model: Model sub-type (e.g., linear regression, generalized additive models, neural networks, etc.)
b. If process-based model: Model sub-type (numerical or analytical), model name (if available), and which processes were represented within the model
4. **Modeling data**: Modeling endpoint, input data, monitoring data characteristics (e.g., duration and frequency), and monitoring methods used for the endpoint
5. **Model application**: How the model was used (e.g., hindcasting, forecasting, prediction between observations in time or space, scenario analysis, sensitivity analysis); how many and which variables and processes were identified as the most important predictors by the authors; the perceived, potential, or actual harms from a HAB that

motivated the modeling; and a concise description of what the authors described as key points and lessons learned from the articles.

### 2.4 Model type descriptions

Because the focus of this literature review was on models, we classified each article according to the modeling approach used: process-based or data-driven. Process-based (mechanistic) models use mathematical equations to represent the physical, chemical, or biological processes that link causes to effects, often through mass balance frameworks. These models range from complex numerical simulations that approximate solutions under realistic, variable conditions to simplified analytical models that provide exact solutions under idealized assumptions. Alternatively, data-driven models, including statistical and machine learning approaches, predict outcomes based on observed data without explicitly defining system processes. They range from simple regressions requiring minimal data to complex neural networks capable of capturing nonlinear, spatiotemporal patterns, though the latter demand large datasets and are more challenging to apply in data-sparse environments. Longer descriptions of these model types are available in the Supporting Materials-1.

Articles containing modeling approaches that did not fit neatly into the process-based or data-driven paradigms were classified as “other”. These include articles that combine process-based and data-driven approaches into one modeling framework (sometimes termed “hybrid” models or frameworks). Such modeling approaches may combine predictions from process-based and data-driven models, replace parts of a data-driven model with process-based components, or use outputs of a process-based model as inputs to a data-driven model (Parshotam and Robertson, 2018; Willard et al., 2022).

If an article described more than one model, we extracted the information holistically for the entire paper and noted how many individual models were developed. As such, summaries are in relation to the number of articles, not the number of models.

## 3. Results and Discussion

In total, we critically reviewed 162 articles with publication dates spanning almost 50 years (1975 – 2023). A bibliography containing citations of the 162 articles is available in Supporting Materials-2. Additionally, Gorney et al. (2025) contains the information compiled from each article during the critical reviews and can be used to locate articles of interest. Reference lists for specific topical themes not easily identifiable in Gorney et al. (2025) are provided in Table SM-1 in Supporting Materials-1.

Most lead authors were affiliated with academic institutions (77% of all articles); 22% of lead authors were affiliated with government entities, and the remaining 4% were affiliated with nongovernmental organizations or consulting firms. Some authors had dual affiliations. Notably, one article was written by a high school student (Claudson, 1975). Articles were published in a wide variety of journals, indicating no preferred publication outlets for the river HAB modeling community. There were approximately 73 unique journals names, with many (n=46) represented by just one article. The most common journals were *Ecological Modeling* (18 articles), *Water Research* (12), and *Water* (9). Five articles were peer reviewed government publications (Table SM-1).

In comparison to Rousso et al.’s (2020) review of lakes and reservoirs (122 articles, 1988 – 2019), our review captured 40 more articles and spanned 18 more years. The timespan of our review is heavily influenced by the earliest publication in the dataset (1975); if this article is excluded, our dataset begins in 1986, around the same time as Rousso et al. (2020). The higher number of articles included in our review can be attributed to our inclusion of all freshwater taxa associated with potentially harmful blooms, as well as the timing of our review relative to Rousso et al. (2020). Had we limited our review to cyanobacteria, 52 articles would have been included, spanning the years 1988 to 2023; if we also constrained our set to the same time span as Rousso et al. (2020), the number of articles would have been 37. The limited number of cyanobacteria-related river HAB modeling articles compared to lakes reflects the propensity for cyanobacteria to dominate in quiescent rather than flowing waters (Chételat et al., 2006; Reynolds and Descy, 1996) and the relative lack of river-focused cyanobacteria studies overall in the literature (Graham et al., 2020). However, the number of articles in our review indicates that algal dynamics and algal-related harms in river systems have been a sustained, and growing, concern for at least half a century.

### 3.1 Environmental setting

Representing over 80 different rivers worldwide, the models developed in the articles were clustered by continent (Figure 2) and country (Table SM-3). The geographic distribution of river HAB modeling articles was generally similar to that observed for lakes and reservoirs (Rousso et al., 2020). Seven percent (n=11) of articles applied models to idealized settings. Of the remaining 151 articles, a majority developed HAB models for river systems in Asia, followed by North America and Europe (Table 2). By country, 26% of the articles were for river systems in South Korea, followed by 21% in the United States and 12% in China (Table SM-3).

**Figure 2.**
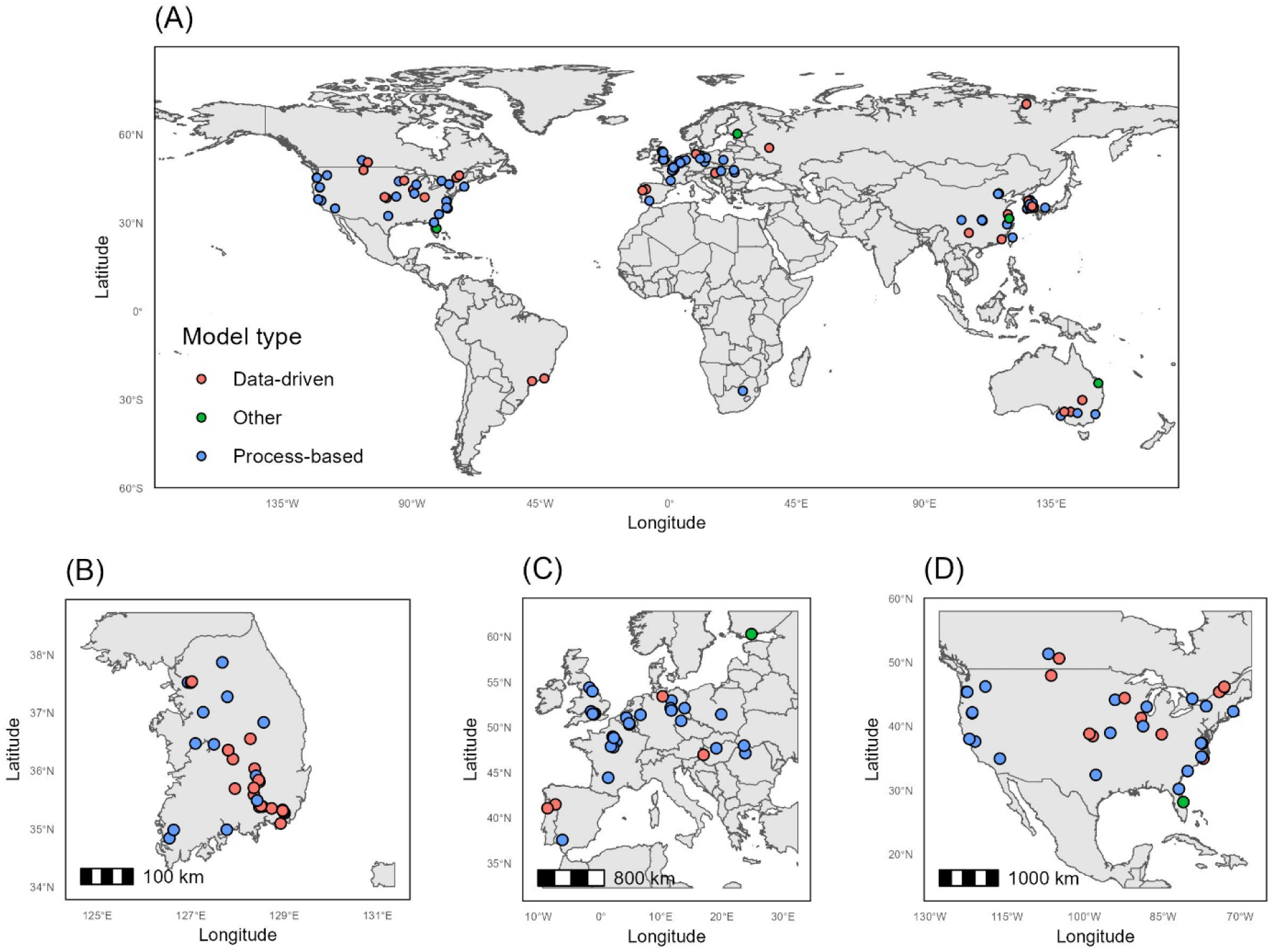
World map (A) of river systems identified in the articles. Map includes only articles that developed models for real systems (151 of 162 articles). Inlays highlight B = South Korea, C = Europe, and D = North America.

**Table 2.**
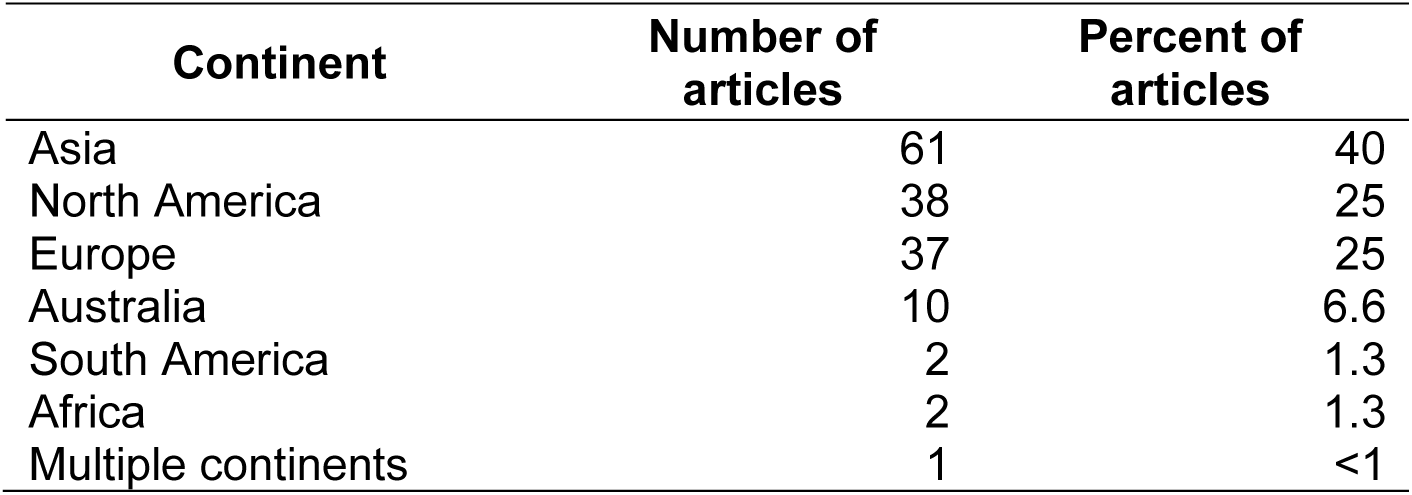
Number and percent of articles that developed models for real systems (n = 151 of 162 articles) across continents.

| Continent | Number of articles | Percent of articles |
| --- | --- | --- |
| Asia | 61 | 40 |
| North America | 38 | 25 |
| Europe | 37 | 25 |
| Australia | 10 | 6.6 |
| South America | 2 | 1.3 |
| Africa | 2 | 1.3 |
| Multiple continents | 1 | <1 |

South Korea was a geographic hotspot for river HABs modeling (Figure 2B), and the Nakdong River–included in 19% of all articles–was a focal point. Furthermore, the top four most common study locations in the literature review are from the four major river basins in South Korea: the Nakdong (n = 29), Han (n = 9), Yeongsan (n = 9), and Geum (n = 8) Rivers. These rivers serve as major drinking-water sources for urban centers of South Korea and are used for industrial and agricultural purposes (Srivastava et al., 2015); as such, the modeling emphasis in these systems underscores their societal importance.

The remaining system-specific articles were more widely distributed geographically and represented over 75 different river systems. Two articles did not report the name of the system being modeled (Crossman et al., 2021; Wang et al., 2019a), and another two articles modeled conditions across multiple systems (Lucas et al., 2009b; Savoy and Harvey, 2023). About a third of the system-specific articles (35%) developed HAB models for one of 53 individual systems. Excluding these articles that modeled unique systems, the top four South Korean river systems and the four articles that did not report a system name or modeled multiple systems leaves about 40% of the system-specific articles (n = 64) that modeled the same system as other researchers. These articles are spread across 23 river systems with between about 2 and 5 articles published per system. As observed by Rousso et al. (2020), few modeling efforts occurred in South America or Africa (n = 2 each), despite widespread and increasing HAB occurrence on these continents (Feng et al., 2024). Both studies in South America were located in Brazil – one on a reservoir (de Souza Beghelli et al., 2016), and one at the catchment scale (Neres-Lima et al., 2017). The Vaal River in South Africa was the focus of the two studies conducted in Africa (Cloot and Roux, 1997; Cloot and Piererse, 1999). Feng et al. (2024) noted that most HAB studies occur in high-income countries, even though low-income areas may face higher HAB occurrence and risk. Therefore, the distribution of river HAB modeling efforts is likely associated with resources available for monitoring, research, and investment in management, as opposed to HAB occurrence and risk.

Non-wadable rivers were the overwhelming focus of river HAB modeling efforts (91% of studies in real settings), most of which were likely large enough to support barge traffic (64%). About half of the articles (47%) describe rivers with in-stream obstructions such as weirs, run-of-river reservoirs, or locks and dams. Given that we included run-of-river reservoirs, there was the potential for articles to be included in both our review and that of Rousso et al. (2020). However, only one paper (focused on the Douro River, Portugal) was captured by both reviews (Teles et al., 2006). About a third of articles focused on free-flowing rivers that did not have nearby in-stream obstructions (31%; Figure 3A). Large rivers with in-stream obstructions were the most frequently modeled systems (41% of all articles), including the four major rivers in South Korea, the Thames River in the United Kingdom, the Xiangxi River in China, and the Seine River in France. Relatively few studies developed models across a river network or watershed (15%) or focused on rivers in connection with lakes (11%; Figure 3A). Similarly, rivers in estuarine settings (n = 21, Table SM-1) draining to the ocean or noted for having tidal influences were the focus of about 13% (Figure 3A) or 9% (Figure 3B) of the articles, respectively.

**Figure 3.**
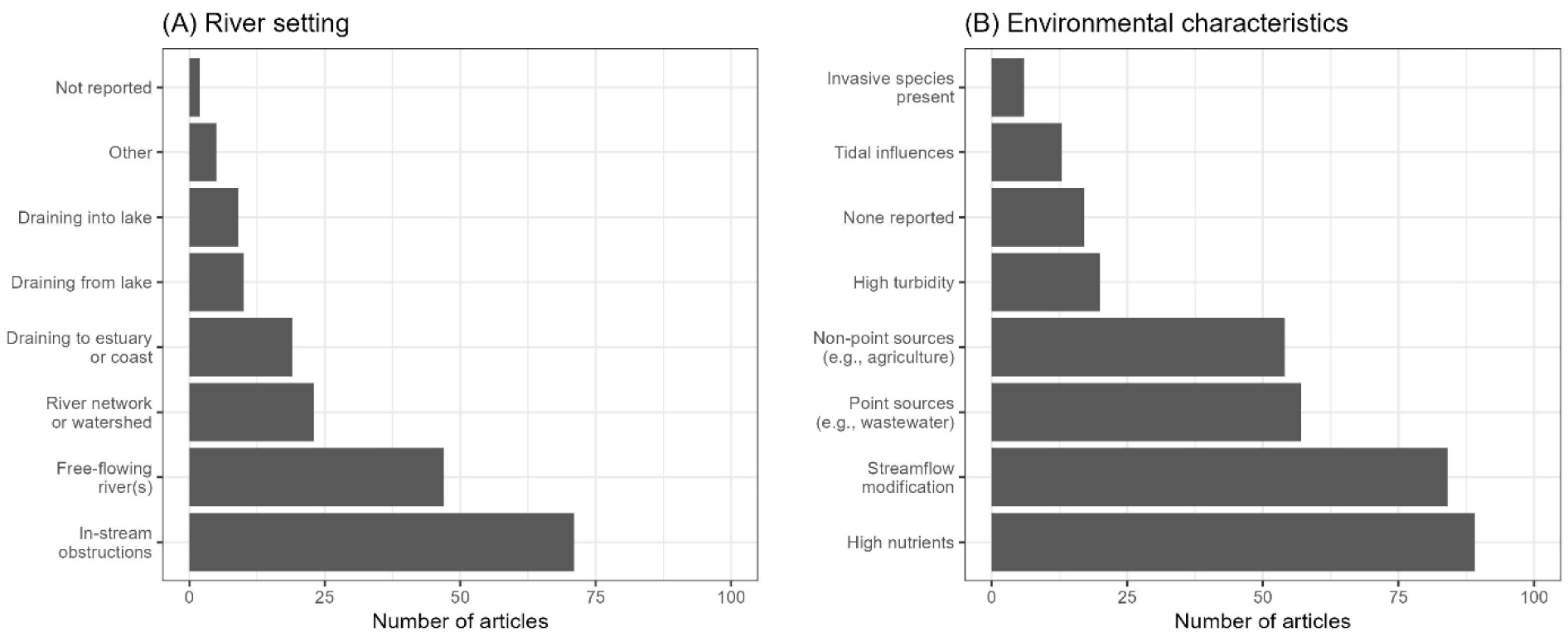
River setting (A) and environmental characteristics of the river setting or watershed described by the article’s author(s) (B) reported in system-specific articles. Percentages will not sum to 100% because an article may model a system that includes multiple river settings (e.g., reaches that are free-flowing and drain from a lake), and often authors mentioned multiple environmental characteristics when describing the riverine setting of their modeling effort. Point and nonpoint sources in (B) refer only to nutrients.

Authors included a variety of characteristics when describing the environmental setting of the modeled system (Figure 3B), many of which indicated degraded or intensively managed systems. Over half the articles mentioned streamflow modification (56%) in terms of diversions, pumping, or other human activities that control the flow rate or volume of water (Figure 3B) which may or may not be related to weirs, dams, or other in-stream obstructions (the latter presented in Figure 3A). Similarly, 59% of articles described eutrophic conditions. Point source (e.g., wastewater discharges) and nonpoint source (e.g., agricultural runoff) influences were described in 38% and 36% of the articles, respectively. Like Rousso et al. (2020), most articles included in our review focused on modeling nutrient enriched systems. Although nutrients are a consistent driver of algal biomass standing crop and HAB formation (Chorus and Welker, 2021; Patiño et al., 2023), nutrient influence in lotic systems is further moderated by streamflow and transport processes that may disconnect HAB occurrence from source conditions (Giblin and Gerrish, 2020; Giblin et al., 2022; Graham et al., 2012; Otten et al., 2015; Schmadel et al., 2024; Williamson et al., 2018; Wood et al., 2020). As such, there would be benefits to additional modeling in systems with complex hydrologic characteristics and geomorphic conditions where causal factors of HABs may not be apparent within the mainstem of the river.

### 3.2 Modeling data

#### 3.2.1 Modeling endpoints

Outputs from the models were diverse. Although most of the articles used algal biomass or a proxy like chlorophyll as the modeling endpoint, a quarter of the reviewed articles (n=42, 26%) used endpoints classified as “other”. Additional endpoints included primary production (n = 5), novel proxies based on oxygen data or metabolism estimates (e.g., Harvey et al., 2024; Wang et al., 2019b), algal cellular nutrient content (n = 2; Bucci et al., 2011; Thebault and Qotbi, 1999), concentrations of taste and odor compounds (n = 1; Chung et al., 2016), probabilities of bloom occurrence (e.g., Kim et al., 2022c; Kim et al., 2021a; Nietch et al., 2022), categorical assignments (e.g., Park et al., 2021), growth rates (Descy et al., 1987; Pinckney et al., 1997), or latent variables (Arhonditsis et al., 2007a; Arhonditsis et al., 2007b), among others. In some articles, models used endpoints that were directly linked to potential harms, such as cyanotoxins (n=8; Gorney et al., 2025). Other articles related model outputs to a threshold or action level. Process-based models often provided estimates of water quality (e.g., dissolved oxygen or nutrient concentrations) and streamflow in addition to an algal-related endpoint. Articles that used data-driven modeling techniques often included multiple models that used either the same modeling endpoint to identify the optimal model formulation or used multiple modeling endpoints to capture various HAB indicators (e.g., cyanobacteria concentration and probability of exceeding a threshold).

When we only considered endpoints that quantified the amount of algae, the presence of certain algal taxa, or both, we encountered a mix of units, analytical methods, and taxonomic levels. These inconsistencies complicated synthesis across the articles—an issue described in detail by Ho and Michalak (2015) for western Lake Erie, USA. About half of the articles (46%, n=75) used an endpoint that quantified the amount of algae present; however, these endpoints included biomass, biovolume, or cell abundance, typically in units of micrograms per liter (µg/L), cubic micrometers per liter (µm^3^/L), or cells per liter (cell/L), respectively. Additionally, these 75 articles used a mix of microscopy (n=41, 55%) and laboratory-based pigment analysis (n=32, 43%) to determine algal quantity endpoints. Some articles used both methods as a means of supplementing the other or to determine taxonomic information. Across all the articles, it was common for the modeling endpoints to indicate the algal community composition, usually at the phylum level or for a particular taxon (41%, n=67). Cyanobacteria were the most common taxa-specific endpoint (32%, n = 52), with genera such as *Microcystis* and *Dolichospermum* (formerly *Anabaena*; throughout this manuscript we refer to this taxon using the name used in the originating article) occurring in multiple articles. Notably, these taxa were also the most frequently modeled in lake systems (Rousso et al., 2020), highlighting the ubiquity of these organisms regardless of hydrologic regime. Other than cyanobacteria, green algae (chlorophytes) and diatoms occurred in 23% of the articles (n=37). For articles that did not provide taxonomic information (n=95, 59%), the modeling endpoint often represented chlorophyll or total phytoplankton.

Chlorophyll was a widely used modeling endpoint across the articles (56% of articles, n = 90), in spite of a long-debated history over analytical methods and use as an indicator of algal biomass in aquatic systems (Schurmann et al., 2024). Some of the ambiguity around the use of chlorophyll is highlighted in previous studies that indicate chlorophyll *a* content per cell (or per unit of biomass) is not consistent across taxa and varies in response to cell physiology and environmental conditions (Cloern et al., 1995; Foster et al., 2022). Additionally, *in situ* measurements via fluorescence sensors are often not directly comparable to extracted laboratory values for a variety of reasons, including light history, cell morphology, nonphotochemical quenching, and water column turbidity (Foster et al., 2022). We did not distinguish between chlorophyll *a* and other chlorophylls in our review and found chlorophyll data were used in a variety of ways. Many articles used chlorophyll concentration (expressed as µg/L) directly as a modeling endpoint (e.g., He et al., 2020; Li et al., 2012; Scharfe et al., 2009; Su et al., 2022), whereas other articles converted measurements like biovolume (e.g., Thebault and Qotbi, 1999) or carbonaceous biomass (e.g., Cerco et al., 2004) to chlorophyll using paired observational data or literature values, and still others proceeded in the reverse direction, converting chlorophyll *a* to other values like phytoplankton biomass. Thebault and Qotbi (1999) compared the influence of using biomass proxies like chlorophyll and total biovolume in a process-based model for the Lot River, France, and found inconsistencies in model output between these measures but generally comparable temporal patterns. Chlorophyll data will likely remain a common modeling endpoint because they are easier and less costly to measure compared to microscopy methods necessary for determining biovolume, cell counts, or taxonomic composition. Chlorophyll was similarly a common endpoint in lake-focused HAB models, as it was used in 69% of the articles in Rousso et al. (2020). A small subset of lake-focused articles used phycocyanin as a modeling endpoint (6%; Rousso et al., 2020); however, phycocyanin was not used in any of the river-focused articles in our literature review, even though cyanobacteria abundance is more closely tied to phycocyanin than chlorophyll in many freshwater systems (Chorus and Welker, 2021).

Out of the 162 articles, algal toxin concentrations—a key HAB concern—were only used as the modeling endpoint for eight articles (8 of 162 or 5%). Of these eight articles, five of the articles used an idealized setting to develop models. These five articles focused on understanding the transport and growth of *Prymnesium parvum* (golden algae) and associated toxins in riverine systems that have stagnant side- and back-channel areas. Grover et al. (2011) derived the first mathematical model that described algal toxin dispersion through this complex riverine setting. This research was extended by the four other articles and in Grover et al. (2017). Hsu et al. (2013) incorporated the influence of zooplankton and their ability to suppress algal abundance and limit toxin concentrations; Abbas (2015) incorporated longitudinal transport and biochemical reaction kinetics into the flowing main-channel portion of the model; Wang (2015) incorporated seasonality; and Wang et al. (2015) considered the role of a limiting nutrient, namely nitrogen. Ultimately, this set of articles identified a reproduction ratio for *P*. *parvum* that differentiated between a washout state and a persistence state in complex riverine systems. The other three articles focusing on algal toxins as a modeling endpoint used field data in data-driven or “other” models to predict concentrations of the cyanotoxin microcystin either spatially or temporally in real river settings (He et al., 2021a; He et al., 2021b; Shan et al., 2022).

Some articles used endpoints in relation to a quantitative threshold or directly predicted a potentially harmful state. For example, two South Korean articles developed models that linked estimates of cell density to algal alert thresholds provided by the government (Kim et al., 2021b; Park et al., 2021). The national South Korean algal alert system was developed in 1997 and focuses on cyanobacterial-related harms (Srivastava et al., 2015). Kim et al. (2021a) studied the four major rivers of South Korea (Han, Nakdong, Geum, and Yeongsan) and modeled the probability of exceeding a single total cyanobacterial abundance threshold of 1,000 cell/mL. Park et al. (2021) studied a single location on the Nakdong River and developed two separate data-driven models to predict the occurrence of conditions related to four “algal alert” categories (normal, caution, warning, and bloom) based on three cell density thresholds (1,000, 10,000, and 1 million cells/mL). He et al. (2021b) provided another example of using a threshold to indicate harm for a collection of sites in eutrophic urban rivers; focusing on the Binhu River Network near Taihu Lake, China, those authors used a classification model that predicts the probability of exceeding a microcystin threshold value of 1.0 µg/L. A model developed by Nietch et al. (2022) for the Ohio River, USA, did not use a quantitative threshold, due to the lack of available data, and instead used a binary endpoint (1 = bloom, 0 = no bloom) based on observational reports of discolored water which were verified to be toxic. Other articles provided semi-quantitative approaches for endpoints that indicate a potentially harmful state. For example, with water suppliers in mind, Rose et al. (2019) developed a risk matrix method to predict phytoplankton-based hazards related to treatment (clogging), aesthetics (taste and odor), and health (cyanotoxins), based on the proximity of the hazard to the water treatment facility and the severity of the consequence. Alternatively, prior to modeling, Hou et al. (2022) used water temperature and bioavailable nutrient concentration data to define five categories ranging from “Potential HAB” to “No potential HAB” and used output from a process-based model to predict the probability of these categories to occur under different simulated scenarios. The diversity of the above examples demonstrates the lack of universal thresholds for recreational and drinking water globally (Brooks et al., 2016; Chorus and Welker, 2021) and the varying ways authors define a HAB. Most articles in the literature review did not use a quantitative threshold or articulate a clear definition of a HAB (*sensu* Gorney et al., 2023). We estimate that less than a quarter of the articles attempted to model a potentially harmful state. It was much more common for models to predict a continuous endpoint, such as chlorophyll *a* concentration, without specifying when that endpoint might be indicative of potential harms.

#### 3.2.2 Input variables

The most common model input variables were nutrients, specifically nitrogen (N) and phosphorus (P), and streamflow (or velocity), used in 72% and 70% of the articles, respectively (Figure 4A). About half the articles used input variables such as

- Water temperature;
- Other hydrologic variables, such as stage, stratification or vertical mixing information, and derived metrics like residence time, flushing rate, or water age;
- Water quality measures other than N and P, such as silica, pH, dissolved oxygen, specific conductance, among others; or
- Light availability, which includes measures of water clarity like turbidity, suspended sediment concentrations, and Secchi depth, in addition to measures like photosynthetically active radiation and irradiance.

**Figure 4.**
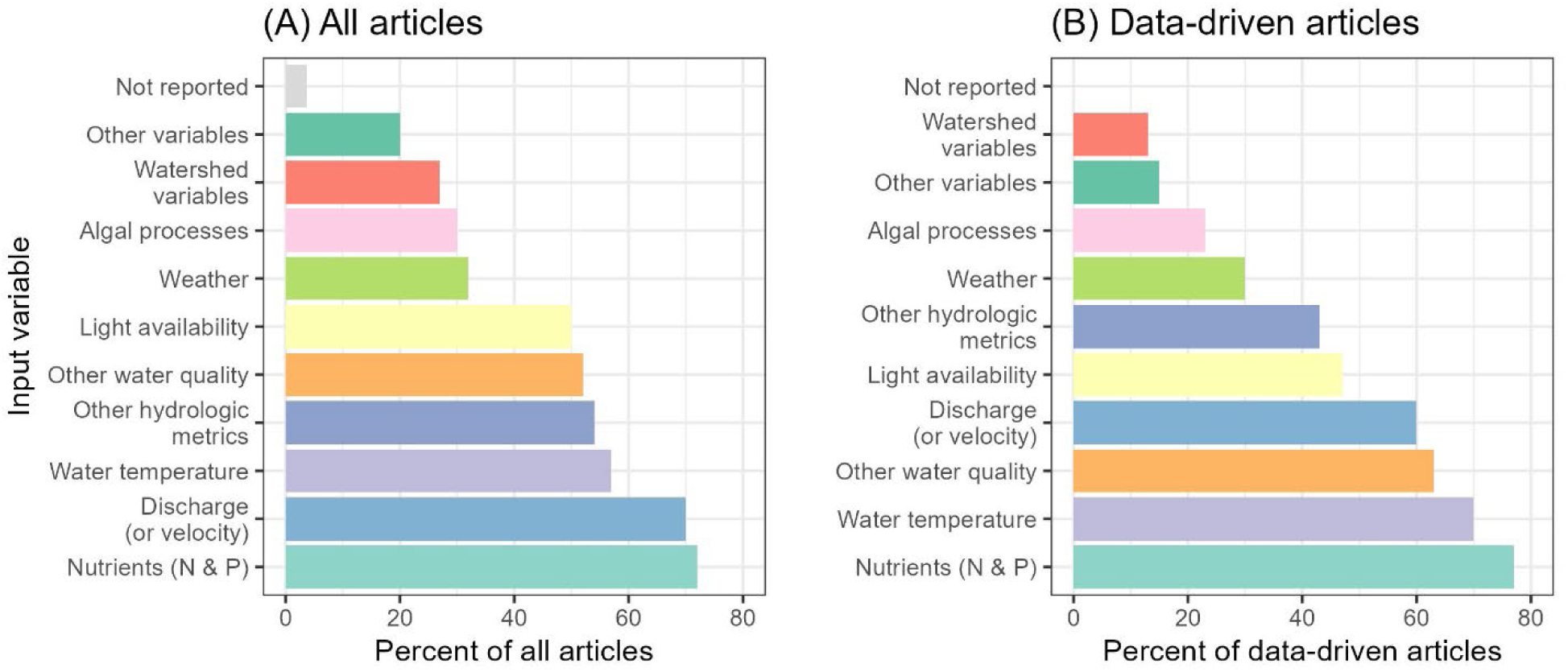
Model input variables indicated across articles, as a percent of (A) all articles and (B) data-driven articles only. Input variables are ordered from least to most common, and colors denote specific input variables. Note: percentages in each panel will not add up to 100 because most models used multiple input variables. [N, nitrogen; P, phosphorus]

Finally, around 30% of the articles used input variables that represented

- weather conditions, such as precipitation, air temperature, wind speed, or wind direction;
- algal processes such algal physiology (e.g., cell sedimentation, buoyancy, motility, resting stages), grazing pressure, and other biologically/ecologically relevant information; or
- watershed characteristics, such as land use characteristics, connections to source areas, among others.

Although subtle, algal processes are likely responsible for important shifts in community composition such as taxonomic succession within broad phytoplankton groups or the dominance of toxigenic strains. However, these types of data are often expensive and difficult to collect, which may partly explain their infrequent use. A small percentage of the articles (less than 10%) did not specify what input variables were used or referred the reader to a different article (Figure 4A).

Rousso et al. (2020) restricted their compilation of input variables to just data-driven models and found water temperature to be the most common input variable followed by pH, light, total phosphorus (TP), and total nitrogen (TN; Table 3). A direct comparison of input variables for data-driven models between lake and river settings is not possible given the different data compilation approaches used by Rousso et al. (2020) and our study; however, broad patterns are discernable (Table 3). Unlike lakes, hydrologic variables were important input variables for river models – 60% of data-driven models included streamflow, velocity, or other hydrologic metrics (Figure 4B); this is not unexpected given that most processes are moderated by streamflow in lotic systems (Cha et al., 2017). Hydrology notwithstanding, nutrients (and other water quality parameters), water temperature, measures of the light environment, and meteorology were the most commonly included input variables in lake and river models, reiterating the importance of these factors in driving algal biomass and community composition (Figure 4B; Table 3). Nutrients were used as input variables more often in river models than lake models, likely because we aggregated all forms of N and P in our literature review, rather than just total nutrients (TN and TP) like Rousso et al. (2020) (Table 3).

**Table 3.** Comparison of the five most common input variables used in lake-focused, data-driven models reported by Rousso et al. (2020) and the river-focused data-driven models of this study. Percentage of river-focused modeling articles using pH is not reported because pH was included in the “other water quality” variables category in our review. Additionally, Rousso et al. (2020) exclusively compiled the percentage of articles that used measures of Secchi depth as an input variable, whereas we included more general measures of light availability such as turbidity, solar radiation, among others.

| **Input variables most commonly reported for data-driven models** | **Percentage of lake-focused data-driven modeling articles (Roussio et al., 2020)** | **Percentage of river-focused data-driven modeling articles (this review)** |
| --- | --- | --- |
| Water temperature | 83% | 70% |
| pH | 58% | --- |
| Phosphorus | 55% (total P) | 67% (all forms of P) |
| Nitrogen | 45% (total N) | 70% (all forms of N) |
| Transparency | 41% (as Secchi depth) | 47% (as “Light availability”) |
| Meteorology | 30% | 30% |

#### 3.2.3 Monitoring data

With the exception of models developed for idealized settings (n = 11), the quality of the calibration data (input variables and modeling endpoint) is important for ensuring appropriate model fit, evaluating model quality, and producing reliable, defensible predictions. As such, for the 151 of 162 articles that developed models for real systems, we summarized the monitoring data in terms of frequency, duration, seasonality, and site counts (Figure 5). We found models were typically developed and calibrated using monitoring data composed of discrete water quality samples collected year-round at 1 to 10 sites, with sampling frequencies ranging from weekly to monthly, over an average duration of five years. However, the distributions of these data characteristics are indicative of a diverse range of data collection regimes (Figure 5). Across the articles, the duration of monitoring ranged from a single year (n = 37) to 35 years (n = 1) and was highly skewed with half the articles using 3-years of data or less (Figure 5A). Similar skew occurred for the number of sites monitored (Figure 5B). The number of monitored sites per article ranged from 1 to 180 sites, and around 75% of the articles used 10 sites or less. In terms of seasonal representativeness, the majority of articles (60%) used monitoring data that were collected year-round. The remaining articles (if seasonality of monitoring was reported), sampled 1 to 8 months of the year (Figure 5C), presumably focused during the HABs season or during HAB events. Most of the articles (68%) used moderately dense datasets with weekly to monthly collection frequencies, likely reflecting the time and effort required to acquire discrete water samples. Lower sampling frequencies were not common. Datasets with daily or sub-daily frequencies were used in the modeling efforts of 26 articles (17% of 151 articles modeling real systems). Many articles used a combination of monitoring frequencies and durations; 12 articles paired daily or sub-daily data collected via *in situ* sensors with data provided via laboratory analysis from discrete water samples (Table SM-1). Finally, 31 articles did not report at least one of the data characteristics presented in Figure 5 and five of these articles did not report information for any of the four data characteristics or referred the reader to another article.

**Figure 5.**
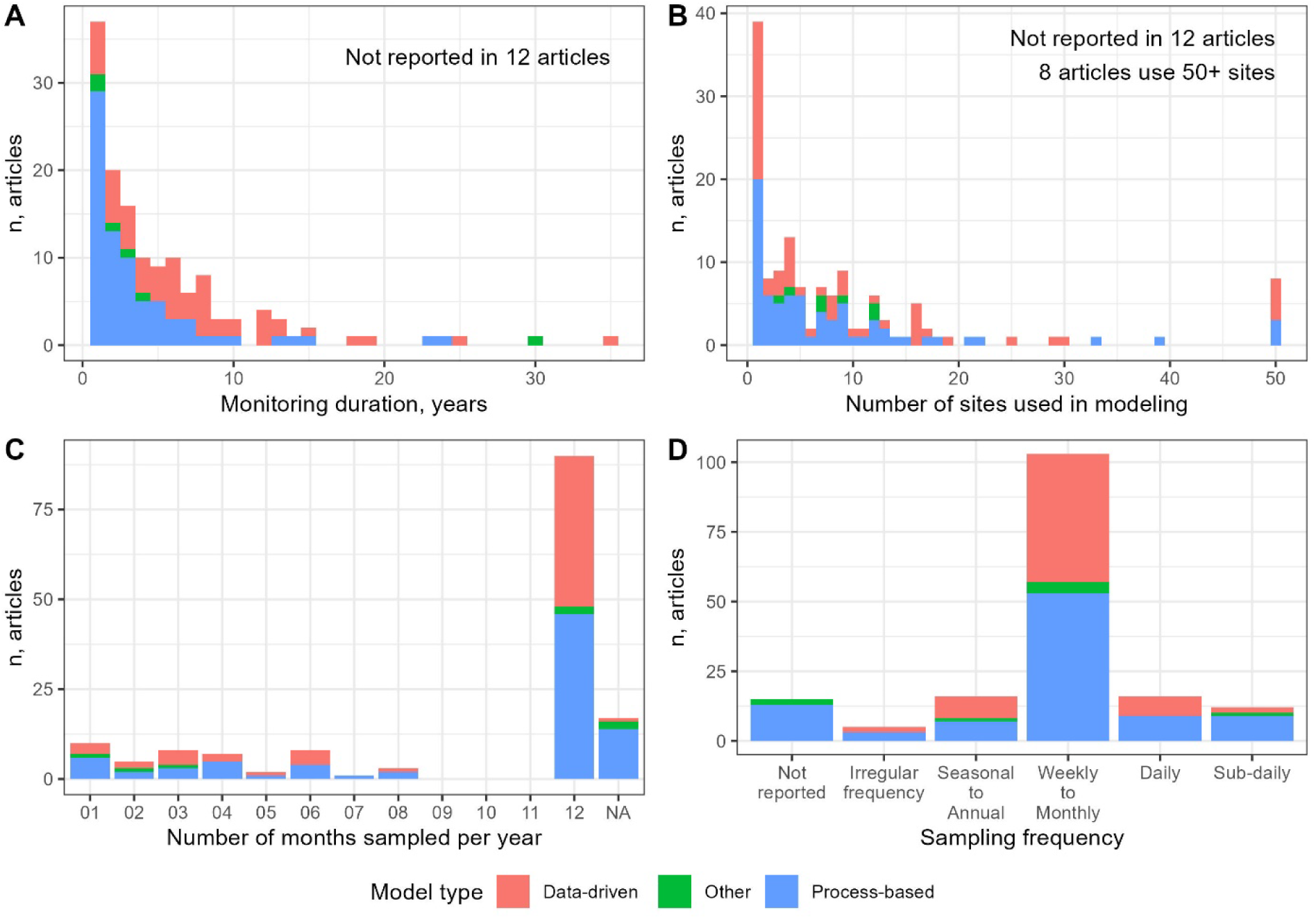
Summary of monitoring data used in articles developing models for real systems (n = 151) in terms of (A) sampling duration in years, (B) number of sites sampled, (C) number of months per year sampled, and (D) sampling frequency. Colors represent model type (data driven, process based, and other). Site counts for the 8 articles that used 50 or more sites (50, 82, 88, 90, 103, 113, 117 and 180) are plotted at x=50.

The use of daily or sub-daily datasets for modeling river HABs has increased over time, starting with 2 articles in 1994-2003 and rising to 9 articles in 2004-2014 and 15 articles during the last decade of our literature review (2014-2023). The two earliest articles were published in 1997. One of the two earliest articles used one year of daily chlorophyll concentrations within a multi-species process-based model and found water temperature, light, and N:P ratio were the drivers of algal growth in the Vaal River, South Africa (Cloot and Roux, 1997). The other article used a variable monitoring frequency to model stratification dynamics of *Anabaena* and *Melosira* (a diatom) behind a river weir in the Murrumbidgee River, Australia. The study included samples collected less than weekly throughout the year, twice weekly during summer months, and sub-daily and at multiple depths during several 24-hours periods (Bormans and Condie, 1998). Similarly, Pathak et al. (2021) used a combination of hourly to weekly water quality information to model diurnal scale phytoplankton dynamics in the Thames River, UK. Many of the high frequency datasets are composed of chlorophyll concentrations based on fluorescence measured by *in situ* sensors. As sensor technology has advanced over the last two decades, deploying and maintaining algal fluorometers has become increasingly common, though still not without challenges (Foster et al., 2022).

We came across several notable datasets during our literature review. Savoy and Harvey (2023) compiled a dataset of daily chlorophyll and explanatory variables for a diverse set of 82 river sites across the U.S. Shan et al. (2022) collected high frequency data using buoy-mounted systems that measured algal cell counts and microcystin concentrations daily, along with a host of additional water quality and physical parameters, in 4 tributaries to the Yangtze River, China. Perhaps one of the most impressive datasets is South Korea’s Water Environment Information System (http://water.nier.go.kr/web) operated by the National Institute of Environmental Research. This dataset contains weekly cell counts for multiple potential cyanotoxin-producing taxa as well as other water quality measurements for multiple river locations across South Korea. This database was used by multiple articles in our literature review, including (Kim et al., 2023; Kim et al., 2022a; Kim et al., 2022b; Kim et al., 2022c; Kim et al., 2022d; Pyo et al., 2019).

### 3.3 Model types

Across the 162 articles, process-based models were more common than data-driven models, appearing in 59% and 37% of the articles, respectively. We classified the remaining 4% (n = 7) of articles as “other” (Guven and Howard, 2007; He et al., 2021a; Pathak et al., 2021; Rankinen et al., 2019; Rose et al., 2019; Yan et al., 2021; Table SM-1), which includes hybrid modeling approaches. In contrast, in lake settings Rousso et al. (2020) found that 40% of articles used process-based models and 60% used data-driven models. Yet, similar to lake models (Rousso et al., 2020), the proportion of data-driven river HAB models has increased over time (Figure 6A). Since the mid-1990s, articles using data-driven models have increased, with a sharp uptick starting in 2004-2013. Process-based models were dominant in the early decades of river HAB modeling but by the most recent decade (2014-2023), data-driven models comprised almost half of articles (Figure 6). Model types also varied geographically (Figure 2). Process-based models were a popular choice in the U.S., especially in coastal areas, and in Europe and South Korea. Data-driven models were most frequently used in South Korea (43% of all data-driven articles) followed by the U.S. and China (Table 2).

**Figure 6.**
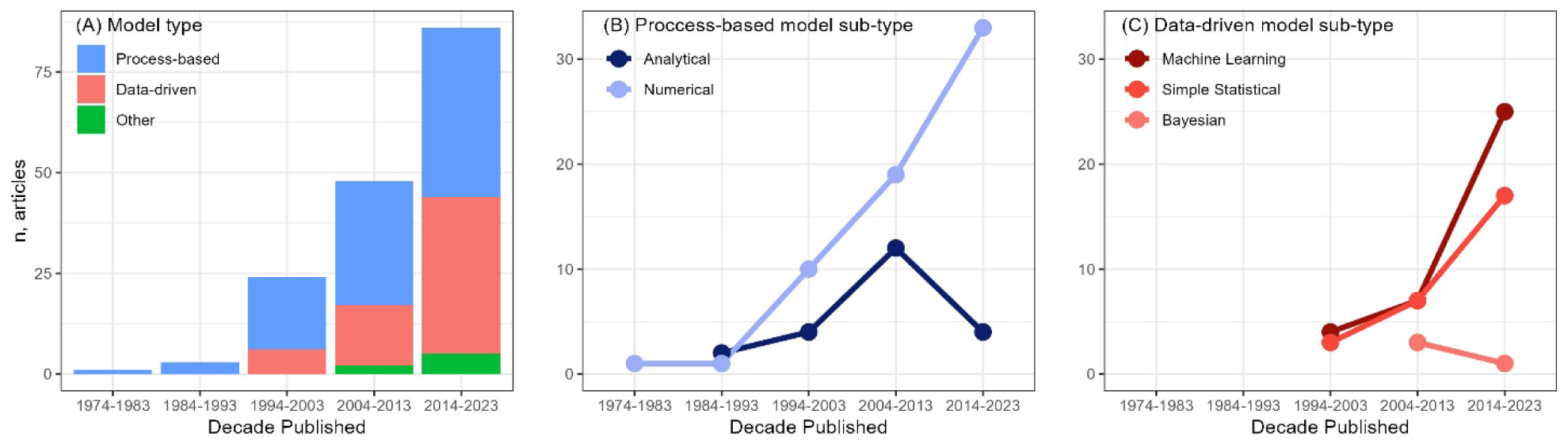
Number of modeling articles published by decade (A) and for major sub-types of process-based (B) and data-driven (C) models. Models are grouped into either numerical or analytical sub-types for process-based models (9 process-based modeling articles that did not report a sub-type are not included) and into machine learning (e.g., neural networks, regression trees), simple statistical (e.g., linear regression, generalized additive models), or Bayesian sub-types for data-driven models. Note difference in y-axis scale between panel (A) and panels (B) and (C).

#### 3.3.1 Process-based models

Of the 95 articles that used only process-based models, 67% were numerical (n=64), 23% were analytical (n=22), and the remaining 9 articles were undiscernible based on our critical review. Numerical models can require considerable computing resources, especially for domains described with significant spatial detail and in multiple dimensions. In contrast, analytical models are typically straightforward and efficient to solve. Within river HABs modeling, there has been a steady increase in the number of numerical models over time (Figure 6). The increase in numerical modeling efforts is likely due to computing advances over the last several decades, as well as burgeoning amounts of observational data with which to drive, calibrate, and validate such models. The intersection between ecohydrology and modeling is notably exemplified by the earliest article in our literature review. Claudson (1975) developed a numerical process-based model for phytoplankton growth in response to chemical and thermal pollution. The publication outlet, *Communications of the Association for Computing Machinery*, reflects novel application of enhanced computing power that was just becoming available for public use at that time.

A wide spectrum of processes was included within the process-based models. For example, a simple first-order loss rate may be specified by the user and included in model (e.g., Engel et al., 2025; Lucas et al., 1999). Alternatively, more complex approaches can be used to dynamically model the zooplankton population and compute grazing or ingestion rates based on the population dynamics (e.g., Wang et al., 2020; Ward et al., 2012). A similar range of computational complexity was apparent across process-based models to represent turbidity, nutrients, hydrodynamics, water temperature, and other influences on algal dynamics. Regardless of how it was represented in a model, we combined all such representations of an individual process to convey which processes were included in some manner in the process-based articles we reviewed (Figure 7). Many processes, including streamflow (or velocity), nutrient availability (N and P), other water quality properties, water temperature, light, and other physical processes (e.g., antecedent streamflow conditions, channel morphology, wind speed and direction), occurred in over 60% and up to 87% of the process-based modeling articles (Figure 7) and were also common input data for many of the data-driven models (Figure 4B).

**Figure 7.**
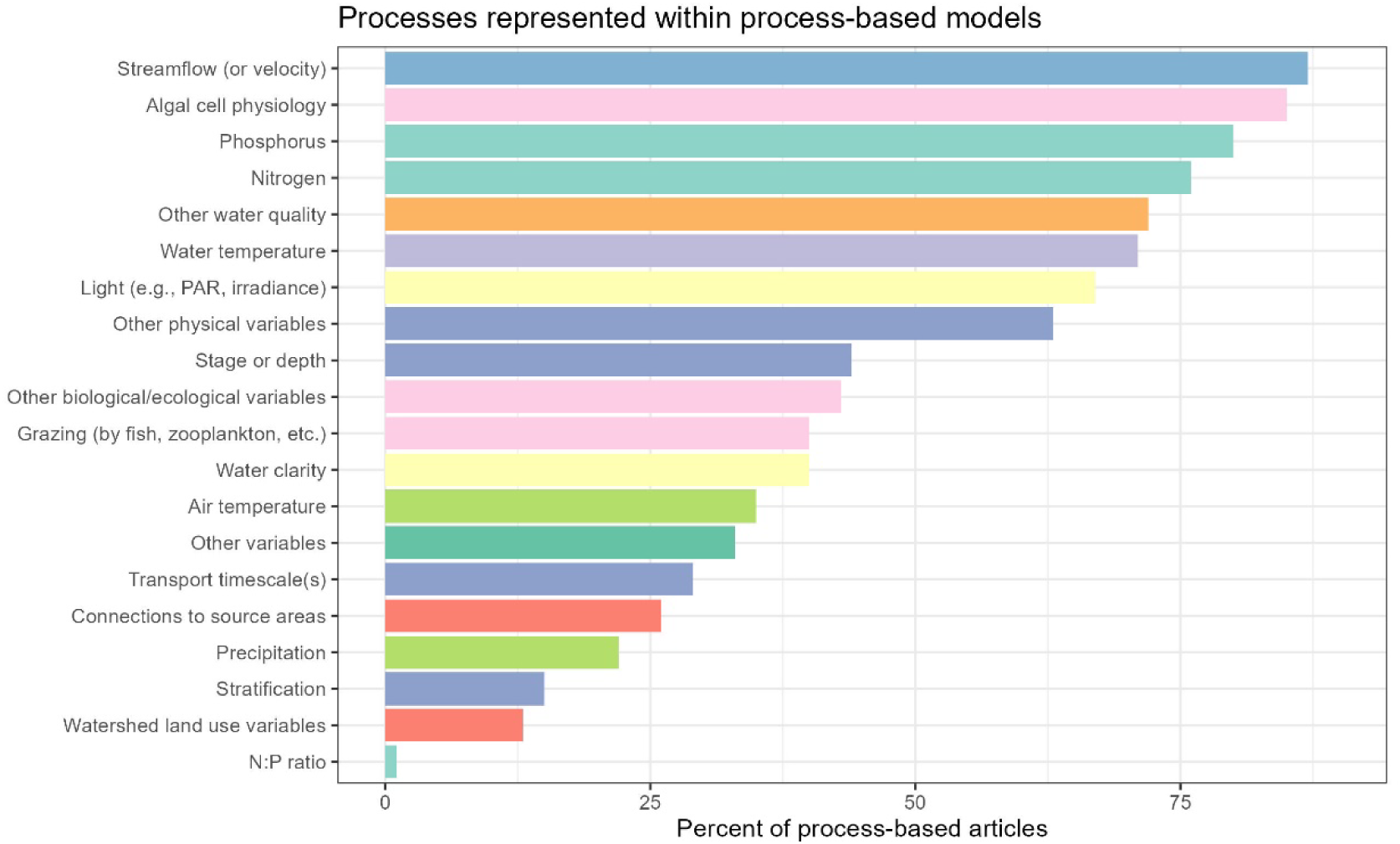
Percent of process-based modeling articles that included a representation of the process listed on the y-axis. Colors correspond to the input variable colors used in Figure 4. The category “other physical variables” includes additional information that describes the physical environment including non-precipitation meteorological data (such as wind direction or speed), antecedent streamflow conditions, channel morphology, among others. [PAR, photosynthetically active radiation; N, nitrogen; P, phosphorus]

A few processes are noteworthy in their comparison to data-driven models or their limited use in process-based models. Algal cell physiology, which included processes such as growth, algal cell sedimentation, buoyancy, or motility, was incorporated into the majority (83%) of process-based models (Figure 7), but when represented more generally as “algal processes”, was much less common in data-driven models (∼22% of data-driven modeling articles). For data-driven models, the “algal processes” input variables in Figure 4B includes algal cell physiology in addition to grazing pressure and other biological/ecological variables. Grazing and connection to source regions or “storage areas” (Grover et al., 2011; Reynolds, 1996; Reynolds and Descy, 1996) are potentially important influences on algal processes represented in 40% and 26% of process-based modeling articles, respectively. Additionally, given that stratification is known in many cases to permit or promote HAB development (Paerl and Huisman, 2008) and was included in 74% of lake HAB models (Rousso et al., 2020), it is somewhat surprising that only 15% of process-based papers (n=14) in our review included that process, as stratified conditions can occur in stagnant side and backwater areas, behind obstructions such as weirs and dams, or during extreme low streamflow conditions. Several of the common process-based models used for rivers (Table 4) can capture stratification processes, if desired. We speculate that the lack of inclusion of stratification in river models is due to a lack of vertical observations in rivers and therefore a lack of evidence about the importance of stratification for HAB development.

**Table 4.** Named process-based models or modeling frameworks implemented in at least three reviewed papers. Model domain: H – hydrology and/or hydrodynamics; WQ – water quality (may include phytoplankton biomass); and E – ecology.

| **Model or Modeling Framework** | **Model domain** | **Number of papers implementing model** | **Reference(s) for model** |
| --- | --- | --- | --- |
| EFDC | H, WQ | 16 | (Hamrick, 1992; U.S. Environmental Protection Agency, 2025) |
| CE-QUAL-W2 | H, WQ, E | 7 | (U.S. Army Corps of Engineers, 2014; Water Quality Research Group, 2025) |
| RIVE/AQUAPHY | WQ, E | 7 | (Gamier et al., 1995) |
| WASP | H, WQ | 7 | (Ambrose, 2017; Wool et al., 2020) |
| RIVERSTRAHLER | H, WQ, E | 4 | (Billen et al., 1994; Gamier et al., 1995) |
| INCA/PERSiST | H, WQ | 3 | (Futter et al., 2014; Whitehead et al., 1998) |
| MIKE | H, WQ, E | 3 | (DHI Group, 2025) |
| PEGASE/<br>POTAMON | H, WQ, E | 3 | (Descy and Gosselain, 1994; Descy et al., 2011; Smitz et al., 1997) |
| QUESTOR | H, WQ | 3 | (Boorman, 2003a; b) |

Many articles used previously published software on which to build process-based models for river HABs (Table 4), and such software may encompass a single model or a framework of models or modules that may be used together. The most common process-based models cited in Rousso et al.’s (2020) study (their Table 1) did not overlap with any in Table 4 herein; this is likely due to differences in dominant governing processes between system types (e.g., horizontal transport in rivers, vertical turbulent mixing in lakes).

#### 3.3.2 Data-driven models

A wide range of techniques was used across the 60 data-driven modeling articles. Similar to findings from Rousso et al. (2020) for lake HAB modeling, most river data-driven modeling articles used a single modeling approach (70%). However, one study compared seven different machine learning models in a river setting (Su et al., 2022). In total, 43 distinct data-driven techniques were identified across all articles (Table SM-4). Artificial neural networks (ANNs) emerged as the most frequently used method, appearing in 18% of the data-driven modeling articles (n = 11). The next most common approach was multiple linear regression, used in 12% of the articles (n = 7), whereas all other techniques were reported in five (8.3%) or fewer articles (Table SM-4). When considering all types of neural networks—such as artificial, recurrent, convolutional, autoencoders, transformers, multilayer perceptron, long short-term memory networks, and others—these methods accounted for 40% of all data-driven modeling articles. Similarly, when aggregating linear regression-based techniques—including simple and multiple linear regression, generalized linear models, structural equation models, mixed effects models, hierarchical models, support vector machines, and others—25% of the data-driven articles employed these methods (Table SM-4). These two categories of techniques were also the two most popular data-driven approaches in the lake HAB modeling review (Rousso et al., 2020). Of the 73 data-driven lake modeling articles in Rousso et al. (2020), 53% used regression techniques (20% used multiple linear regression) and 31% used neural networks (20% used artificial neural networks). Rousso et al. (2020) compiled 28 distinct data-driven techniques across the lake models, many of which we also found in the river modeling articles including logistic regression, support vector machines, recurrent neural network, self-organizing map, evolutionary algorithm (13 lake models used this technique compared to just 1 river model), multiple types of decision trees, multiple types of Bayesian models, and fuzzy logic (Table SM-4). Even amongst the many techniques we identified across the river models, notable lake model techniques missing from our compilation include multiple adaptive regression splines, agent model, elastic network, and Cusp catastrophe theory.

At a broader level, machine learning models (including neural networks and other approaches) were more prevalent than traditional statistical techniques. Although the use of these methods has historically been similar, machine learning techniques have gained prominence over the last decade (Figure 6). Typically, machine learning methods require more data compared to simpler statistical techniques and process-based models. This is due to several factors: machine learning models have a larger number of parameters to estimate, are more susceptible to overfitting, and excel at identifying patterns in complex, interacting variables and non-linear relationships—all of which necessitate larger datasets. Machine learning articles used longer duration monitoring datasets (mean = 7.4 years, median = 6 years) compared to simple statistical (mean = 6.0 years, median = 4 years) or process-based articles (mean = 3.8 years, median = 2 years). Recent increases in environmental data, particularly water quality data (Read et al., 2017), combined with the availability of open-source programming software such as R and Python that supports machine learning modules, have empowered researchers to apply advanced data-driven models to water quality prediction challenges, including riverine HABs modeling. As environmental data volume continues to grow and access to open-source software expands, we anticipate that the upward trend of applying machine learning models will persist.

Despite machine learning models using datasets with slightly longer monitoring durations, they typically drew data from fewer sites compared to simple statistical and process-based models. The median number of sites used by machine learning models was 2.5, compared to 11 sites for simple statistical models and 4.5 sites for process-based models. Neural networks, the most complex data-driven models, perform best when trained on data from a diverse range of sites, allowing them to learn general patterns in hydrological and ecological phenomena (Kratzert et al., 2024). Therefore, it is surprising that machine learning models relied on the fewest number of sites, although our literature review did identify instances of machine learning models trained on a larger number of sites (e.g., 82 sites in Savoy and Harvey, 2023). Only recently, since 2023, have large, curated datasets of lotic algal biomass, such as chlorophyll levels, become available, enabling model training across various site types (Fernandez et al., 2025; Savoy and Harvey, 2023; Spaulding et al., 2024). In contrast, most data-driven applications we reviewed focused on curating datasets for specific, local HABs concerns. However, we propose that by leveraging these recently published large datasets alongside advanced data-driven techniques, prediction accuracy of river HABs can likely improve across a greater number of sites.

As previously mentioned, South Korea, particularly the Nakdong River, is a hotspot for river HAB modeling as 42% (n=25) of all the data-driven modeling articles originated from this location. The collective research efforts in South Korea offer valuable insights into effective data-driven modeling techniques. Among the reviewed articles, 20 of the 25 South Korean data-driven studies used machine learning techniques, often demonstrating superior performance of advanced machine learning methods compared to traditional approaches. For instance, neural networks with temporal awareness, such as recurrent neural networks (RNNs), outperformed simpler statistical methods (Jeong et al., 2008; Lee and Lee, 2018). Furthermore, customized enhancements to RNNs designed to address missing data and low-frequency events showed improved performance over standard RNNs (Kim et al., 2022d). This collection of articles highlighted the strengths of convolutional neural networks (CNNs) in integrating various data types. For example, CNNs effectively incorporated hyperspectral chlorophyll *a* maps, leading to improved predictions compared to the EFDC process-based model (Pyo et al., 2021). Additionally, CNNs were able to use synthetic outputs from an EFDC model to enhance cyanobacterial forecasts (Pyo et al., 2020) which could possibly be considered a hybrid approach, though we classified this article as data-driven. Their ability to leverage spatial information gives CNNs a distinct advantage over other data-driven models. Lee et al. (2022) demonstrated that CNNs could use data from multiple sites to enhance predictions at individual sites within a river network (Lee et al., 2022). Researchers have also recognized the value of simpler statistical methods for predicting river cyanobacterial blooms. For example, (Kim et al., 2020) found that a logistic regression model using just water temperature, river velocity, and phosphorus concentrations achieved over 75% forecast accuracy in South Korean rivers. The application of data-driven methods in select study systems in South Korea illustrates how in-depth research in specific locations can facilitate the comparison and evaluation of various modeling techniques.

#### 3.3.3 Other models

The seven articles containing models we classified as “other” did not fit cleanly into the process-based or data-driven model categories. Some implemented data-driven models to analyze process-based model outputs in order to draw inferences regarding importance of model parameters (Guven and Howard, 2007), relations between various environmental parameters and chlorophyll concentration (Pathak et al., 2021), or lateral bloom position (Yan et al., 2021). Some papers in this category implemented hybrid approaches that combined process-based and data-driven approaches to predict a HAB-related endpoint. For example, He et al. (2021a) used outputs of a process-based hydro-biogeochemical model as inputs to an ANN model to predict microcystin concentrations. Similarly, Rankinen et al. (2019) and Mitrovic et al. (2006) used process-based models to compute environmental conditions (e.g., runoff, streamflows, nutrient concentrations) as inputs to empirical models that predicted, respectively, chlorophyll *a* concentration and bloom frequency. The “risk matrix” approach of Rose et al. (2019) was included in this model category because it represented (relative to the other models in the articles we reviewed) an unusual and “semi-quantitative” approach to assessing cyanobacteria-related risk to a water supply.

#### 3.3.4 Process-based and data-driven model comparisons

Among the 162 articles, 62% reported at least one quantitative measure of model skill. The most used metrics were the coefficient of determination (R²) and root mean square error (RMSE), which appeared in 31% and 25% of the articles, respectively, paralleling findings from Rousso et al. (2020) for lake models. Furthermore, similar to Rousso et al. (2020), we also found a lack of standardization of skill metrics. Across river models, skills metrics categorized as “other” were used in 28% of the articles, making it the second most frequently reported category. Unexpectedly, 38% of the articles did not provide a quantitative measure of model skill, and a substantial proportion of these articles used process-based models. Across the process-based modeling articles, 55% did not report any quantitative metrics, compared to only 12% of data-driven modeling articles (Figure 8). Among the process-based articles that did not report a quantitative skill metric (n = 52), 58% (30 articles) included visual comparisons of observed values and model outputs, either through plots or qualitative descriptions. Model performance metrics enable comparisons across different modeling approaches, driving advancements in both theory and model development (Lewis et al., 2022). Additionally, these metrics effectively capture the insights gained from visual inspections and expert evaluations of model performance (Gauch et al., 2023). Findings from literature reviews of models in river (this study) and lake (Rousso et al., 2020) settings emphasize these points in the context of HABs. The standardization of skill metrics and the use of at least one quantitative metric in future HAB modeling studies could result in better comparisons across different approaches and aid in model selection for river and lake settings.

**Figure 8.**
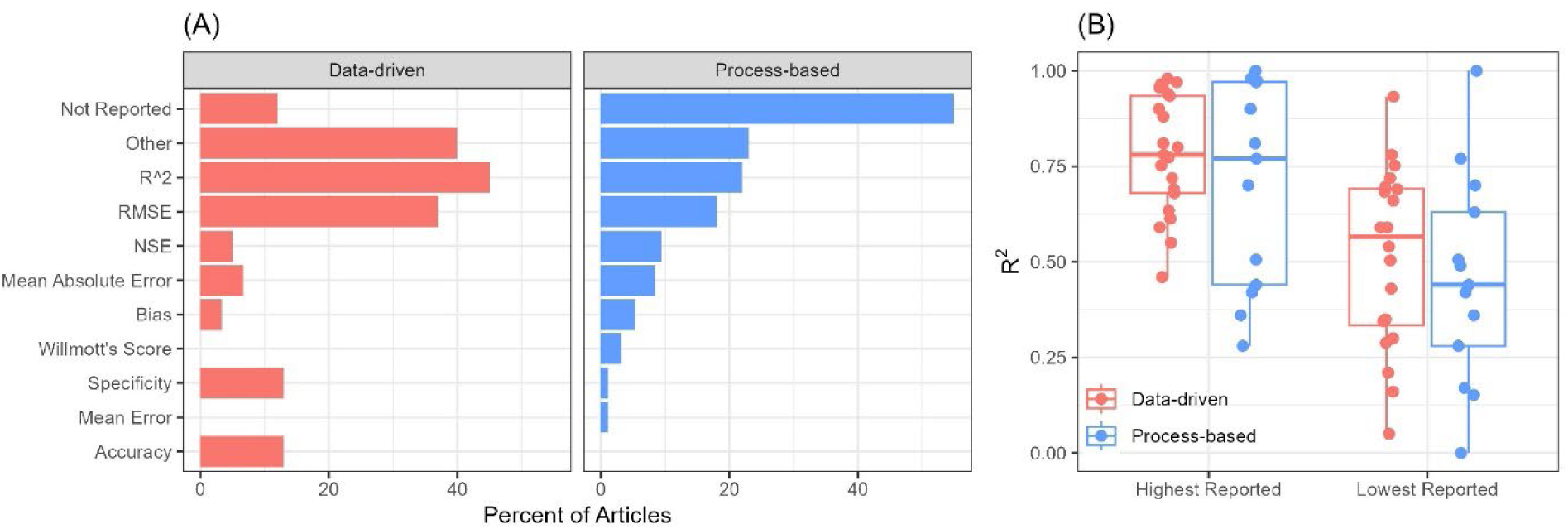
(A) Percentage of articles reporting specific quantitative metrics for assessing model skill, categorized by model type. The quantitative metrics are arranged in order of prevalence among process-based articles. [RMSE, root mean square error; NSE, Nash-Sutcliffe Efficiency] (B) Range of highest and lowest reported R² values from data-driven (n = 21) and process-based (n = 13) modeling articles.

The quantitative metrics demonstrate a range of skill for river HAB modeling. Within individual articles, R² values were reported in various contexts, including testing datasets, training datasets, multiple models, and different endpoints. For each article, we recorded the highest and lowest R² values among all HAB-relevant endpoints, excluding any quantitative model performance metrics related to other physical or chemical outputs. The full range of reported R² values spanned from <0.001 to 0.999, with little variation based on the type of model used (Figure 8). Notably, the lowest R² values reported in process-based modeling articles were generally lower than those in data-driven modeling studies. However, the median of the highest reported R² values was quite similar between the two modeling types, at 0.77 for process-based models and 0.78 for data-driven models (Figure 8B).

To compare the primary predictors of river HABs between process-based and data-driven models, we compiled what the authors identified as the most important predictors indicated by their modeling efforts (Figure 9), a topic discussed by most authors (n = 150). Allowing the articles’ authors to determine important predictors reconciles the disconnect between input data and simulated process with data-driven and process-based models. Both model types use input data in model training or calibration. However, process-based models also use data in additional ways (e.g., setting boundary or initial conditions) and include simulated processes within the model. Input data and simulated processes both have the potential to indicate important predictors of river HABs and thus compiling author-reported predictors synthesizes these two types of information (Figure 9A). Across these 150 articles, authors identified, on average, three important predictors. Nutrients were identified as the most important predictor (39%, n = 63). Light availability, streamflow (or velocity), and algal processes all ranked 2^nd^ (each at 33%), followed closely by water temperature (30%, Figure 9A). We surmise that broad drivers like nutrients, streamflow, and light are likely decent predictors of overall biomass but may struggle to predict finer scale dynamics like seasonal succession in algal community composition. Rousso et al. (2020) compiled the single most important predictor from each lake-focused article in their review. As a percent of all reviewed lake articles (n = 122) they are:

- Water temperature (31.5%)
- Nutrients (23.5%, includes N and P)
- Metrological variables (12%, includes air temperature, wind speed, air pressure, and rainfall)
- Biological variables (6%)
- Water level (5%)
- Land use (3%)
- Streamflow (2%)
- Other water quality (no percentage given, but provided examples for dissolved oxygen, conductivity, and silica).

**Figure 9.**
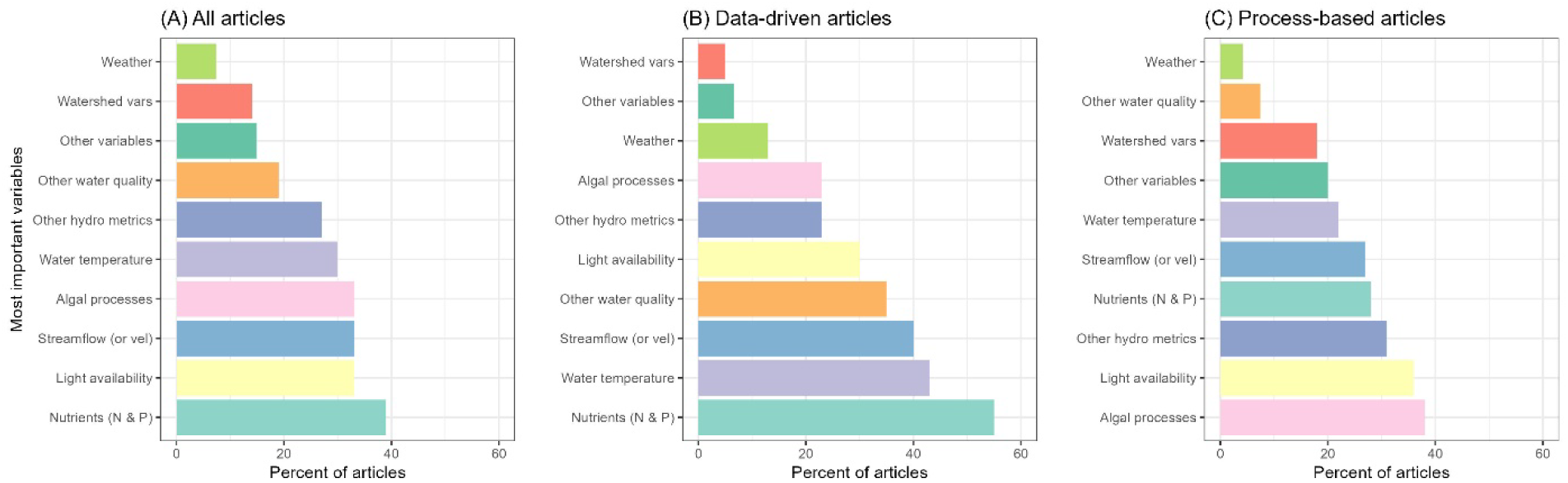
Most important predictors identified by authors during their modeling effort, as a percent of all articles that reported this information (A, n = 150), data-driven modeling articles (B, n = 56), and process-based modeling articles (C, n = 87), ordered from least to most common. Colors denote specific variables and are the same between the panels. Note, the “all articles” plot includes 7 articles with “other” model types. Percentages in each panel will not sum to 100 because most articles reported more than one important predictor. [vel, velocity; N, nitrogen; P, phosphorus]

Three of the five top predictors are shared between lake and river models—nutrients, algal processes/biological variables, and water temperature—despite differences in what was compiled during the two literature reviews (i.e., the single key predictor versus all key predictors from the lake-focused and river-focused literature reviews, respectively; Figure 9A). Not surprisingly, streamflow was ranked as the 3^rd^ most important predictor for river models but ranked 7^th^ for the lake articles (Rousso et al., 2020). However, light availability ranked as the 2^nd^ most important predictor for river models but was never the primary predictor for lake models (Rousso et al., 2020). This discrepancy may be due to differences in how this information was compiled; we used a broad definition of light availability that included measures like photosynthetically active radiation, irradiance, and measures of water clarity (e.g., turbidity, Secchi depth) whereas Rousso et al. (2020) compiled exclusively solar radiation.

When constrained by model type, the primary predictors in river models change substantially (Figure 9). Within the data-driven articles, nutrients, water temperature, streamflow (or velocity), water quality other than nutrients, and light availability remain important predictors in 30-55% of the articles (Figure 9B). For process-based articles, the most important predictor was algal processes (includes predation, cell physiology, and other biological/ecological variables), which was identified as a main predictor in less than a quarter of the data-driven articles. Algal processes were closely followed by light availability and hydrologic metrics (other than streamflow or velocity, e.g., residence time) for process-based models (Figure 9C). Nutrients and streamflow (or velocity) have lower importance in process-based articles compared to data-driven articles, and other hydrologic metrics rise in importance (Figure 9). Some differences in ranking are related to the input data used to drive data-driven models (Figure 4) and the additional processes simulated within process-based models (Figure 7). For example, algal processes are an important feature included in many process-based models, though are rarely included in the input data used to train data-driven models (Figure 4B). Algal processes encompass information like growth rates, nutrient uptake, cell sedimentation, cell motility, and rates of predation. These processes are not readily observed quantities and are rarely, if ever, part of routine or event-based monitoring programs, which data-driven models heavily rely upon. Nevertheless, performance of the best models is roughly comparable between model types (as demonstrated with the median highest-reported R^2^ values in Figure 8). As such, data-driven models may be capturing algal proliferation as a stochastic process, whereas these are deterministic processes within process-based models.

### 3.4 Model application

In over half of the articles (60%), authors articulated multiple purposes for developing or applying a HAB model in a given river setting (Figure 10). The most commonly stated purpose was to make predictions between observations in time (40% of articles), followed by testing different management or climate scenarios (35% of articles), hindcasting (32%), or conducting sensitivity analyses (30%). Both data-driven and process-based models were frequently used to make predictions between observations in time (ranked first or second for model purpose), however the frequency of other purposes varied according to model type (Figure 10).

**Figure 10.**
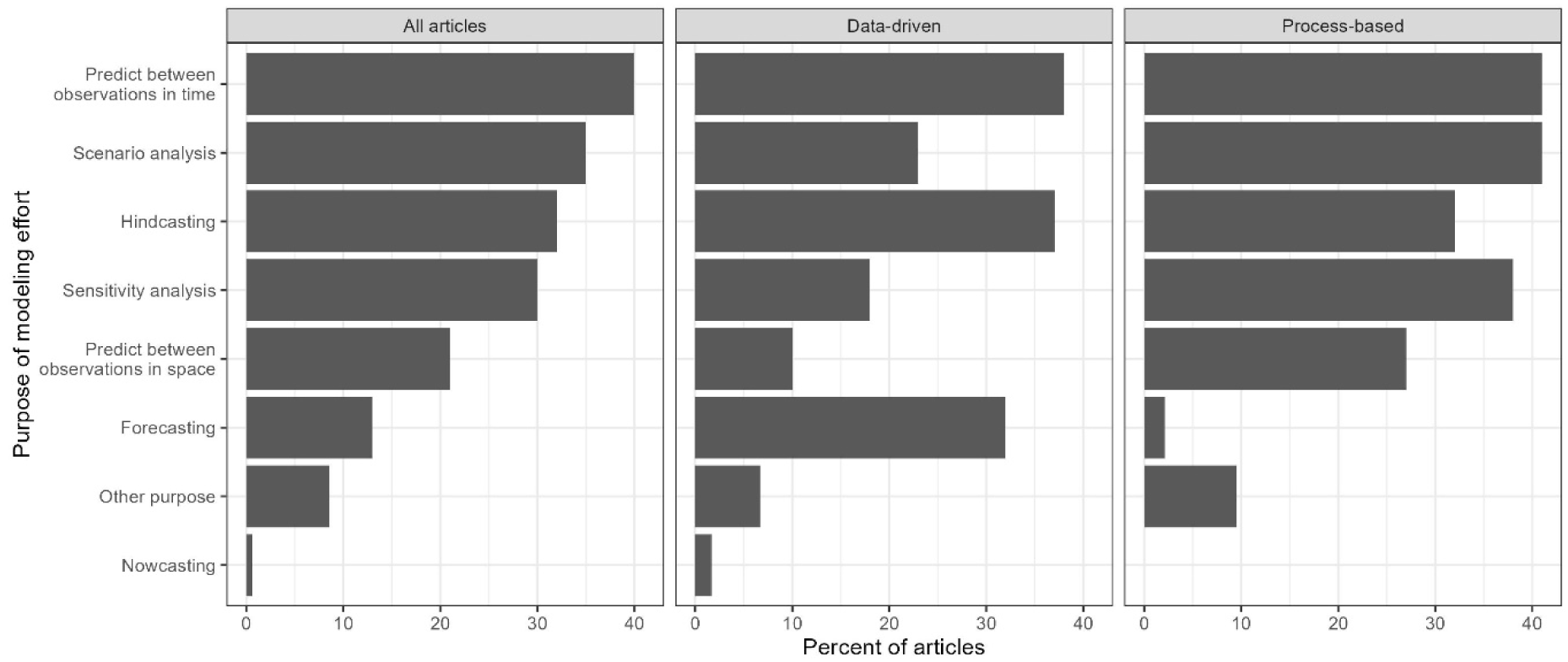
Purpose of river HAB modeling effort as a percent of all articles (n = 162), data-driven modeling articles (n = 60), and process-based modeling articles (n = 95). Note, the “All articles” panel includes seven articles with models classified as “other”.

Scenario analysis was the most common purpose of process-based modeling articles (41% of all process-based modeling articles; tied with predicting between observations in time) yet the 4^th^ most common goal for data-driven models (23% of all data-driven modeling articles; Figure 10). The higher use of process-based models for scenario testing is likely because they incorporate physical laws and accepted process understanding making them more reliable for unobserved conditions. In contrast, data-driven models rely entirely on training data to identify relations between input and output variables and thus may provide less reliable predictions outside of the conditions captured in training datasets (Appling et al., 2022). Considering both process-based and data-driven models (n=57 articles), we identified 17 types of scenarios (Figure SM-1). Authors also explored a variety of scenarios related to changes in climate, key drivers (e.g., moisture conditions, light limitation), the physical environment (e.g., riparian zone characteristics, stratification, land use change), and seasonality. The most common scenarios explored were those related to potential long-term streamflow and nutrient management solutions. The effects of streamflow modification or nutrient reduction strategies on point and nonpoint sources were evaluated in 35% and 28% of these articles, respectively (Figure SM-1), reiterating the influences of these drivers on accumulation of algal biomass in lotic systems and the ability for managers to manipulate these inputs (as opposed to climate effects) to minimizing HAB occurrence.

Similar to our review, Rousso et al. (2020) also found many articles explored longer term mitigations strategies particularly for nutrient management. Articles that explored scenarios for short-term mitigations once a HAB was already present were largely absent from the lake modeling literature. For river models, short-term mitigation scenarios mostly focused on the use of increased streamflows, often released from an upstream reservoir and dam, to suppress the formation of a downstream river HAB (e.g., Mitrovic et al., 2006; Yoshioka and Yaegashi, 2017). Only one article in our review modeled direct HAB management via algicide dosing, specifically copper sulfate (Lewis et al., 2002). Despite the wide range of available short-term mitigation options including physical, chemical, and biological approaches for an established HAB (Burford et al., 2019), scenario testing for these strategies is largely absent from the HAB modeling literature. We do not know if these studies are not being done or if they are being explored outside the realm of predictive and forecasting models and thus would have been excluded from our review.

Forecasting, along with hindcasting and predicting between observations in time, was among the three most common goals of data-driven modeling articles (32%, 37% and 38% of all data-driven articles, respectively; Figure 10). However, only two process-based modeling articles produced forecasts (Ahn et al., 2021; Loos et al., 2020). Within data-driven articles, artificial neural networks were the most common modeling approach used for making forecasts (n = 5, Table SM-4). Additional forecasting methods were diverse and included multiple linear regression, Markov chain, kernel density estimation, long short-term memory, support vector machines, among others. The most common forecasted output from these articles included cell abundance or density with an indication of the taxa present. Of all the articles that provided forecasts (n = 21), most are short-term with lead times ranging from 1-day to 30-days ahead (n = 20). The forecast horizon for potentially harmful events is not more than a few days to weeks, given the influence of weather on the physical and chemical environment in aquatic ecosystems (Benincà et al., 2008; Burford et al., 2020; Petchey et al., 2015). The most common forecast horizon across the articles was 7 days (n = 10). One article provided a 55-day ahead forecast (e.g., Kim et al., 2022c; Kim et al., 2021a; Nietch et al., 2022). Some articles provided multiple forecasting windows. Finally, forecasting models were applied primarily in South Korea (n = 14) with a few in Australia (n = 3), and one each in Portugal, United States, Germany and China.

An important consideration when defining a HAB is a recognition of perceived, potential, or actual harm to humans, animals, the environment, or the economy. We selected this set of 162 articles for our literature review because HABs were motivating the modeling effort, as described by the authors in the title, keywords, or abstract (refer to query keywords, Table 1). During review of these articles, we cataloged the perceived, potential or actual harms described by the authors. Although all articles described at least one harm of concern, about half of the articles specified more than one. A broad category of “general ecosystem health”, which includes eutrophication concerns, was indicated as the motivation for HAB modeling in 85% of the articles. The second most frequent concern was negative effects on drinking water (31%), followed by toxicity and toxin production (23%), and events that affect wild and domestic animals (19%). These potential harms reflect the size and environmental characteristics of the river settings described above, meaning many articles developed models for large rivers and run-of-river reservoirs, which are often used as drinking water sources and for recreation. Although concerns about algal toxins motivated model development in 23% of the articles (n=38), only eight articles explicitly modeled algal toxins in river systems (refer to Modeling endpoints section 3.2.1).

Finally, we summarized key lessons from each article, as indicated by the authors (Figure SM-2). Most of the key lessons pertained to the identification of the main processes driving algal proliferation in the environmental setting being modeled. About a third of the articles found that streamflow conditions in terms of hydrodynamics, discharge rate, transport process, water withdrawals, presence of weirs, operation of dams, and variables like water age were key parts of explaining variability in algal abundance and HAB occurrence. Nutrient concentrations and loads were also found to be an important driving factor in 22% of articles. However, only 9 articles described both streamflow and nutrients as co-drivers. Finally, 19% of the articles (n = 31) had key findings about the specific model presented in the article. For example, these articles described how the model was developed, how the model performed, or tested and described the uncertainty associated with the model, as opposed to providing insights into the processes controlling HAB development, duration, or decline.

### 3.5 Limitations of our study

This systematic literature review has several caveats, especially concerning the statistics derived from our final set of 162 articles. As noted in Section 2.3, to collect consistent information from each critically reviewed article, we populated a fillable form designed for this study. However, in many cases, the desired information was either not addressed at all, briefly alluded to, or described with scant detail in the article. In some cases, the articles referred readers to other papers for such details. At times we made inferences and in other cases we were unable to record information for a particular question. Consequently, there is some uncertainty in the compiled information. Areas in which information was at times incomplete, minimal, or completely absent included:

1. Environmental setting: river size and environmental characteristics
2. Monitoring data: number of sites, frequency, duration, and/or methods related to the collection and laboratory analysis of the observational data used in the modeling process
3. Model details (process-based models): processes included and how they were handled in the model (e.g., specification of rate constants versus dynamic computation); solution methods (numerical versus analytical); treatment of time (steady-state versus time-varying)
4. Model input data or parameters (process-based models): description of all model inputs and parameters derived from literature
5. Model calibration and validation: use of calibration versus validation dataset; specifics about the model formulation; descriptions of how models were validated including calculation of quantitative skill metrics

Additionally, we acknowledge the diversity of the scientific questions and modeling goals of these 162 articles and that the findings from different scientific questions may not be directly comparable. For example, we compiled the main predictors identified by the authors; however, what the authors considered a main predictor is highly dependent on the scientific question being addressed and the parameters for which model sensitivity was tested. This caveat applies to many of the questions related to model application, which we attempted to capture using questions about model purpose and key lessons.

Furthermore, even though we followed a systematic approach to this literature review (similar to Rousso et al., 2020, and described by Pickering and Byrne, 2014) there are two places where we may have inadvertently excluded a relevant article. First, was during the development of the query used in Phase 1. To address this concern, we used a set of 23 validation articles containing river HAB models to develop and refine the search query (Table SM-1). Second, was during the three-step screening in Phase 2. Here we used explicit inclusion/exclusion criteria (Table SM-2), but there may have been slight differences in the interpretation of this criteria depending on the reviewer. Finally, our restriction to only English-language articles may be excluding river HAB modeling literature in areas of the world that are not dominated by English speakers.

### 3.6 Challenges to advancing river HAB modeling

Using this collection of river HAB modeling articles, we identified three challenges to advancing river HAB modeling. First, is a lack of available data, especially for key processes, and the difficulty of incorporating newer data streams such as high frequency *in situ* data or remote sensing to models. Second, is the geographic clustering of modeling efforts and their focus on a particular river archetype which neglects important habitats and areas of the world. Third, is the challenge of synthesizing results across various model inputs, model endpoints, and model evaluations metrics presented across the articles which are at times poorly reported or not comparable.

#### 3.6.1 Data gaps

One of the most prominent challenges limiting the ability to model water quality in aquatic systems is the lack of data (Lucas et al., 2025). For process-based water quality modeling generally, “Data related gaps are extensive and include the nearly universal need for improved datasets for model calibration, validation, and specification of boundary conditions; biogeochemical thermodynamic and kinetic rate parameters; constituent sources; and data describing the physical setting…” (Lucas et al., 2025). These general data needs apply at least as aptly to modeling HABs. Rousso et al. (2020) noted a similar challenge for lake HAB models, pointing to constraints imposed by not only the amount but also the quality of available data (including monitoring technologies and data frequencies) and how this influences process-based and data-driven modeling approaches differently.

Similar to the lake-focused review (Rousso et al., 2020), we also found process-based models for river HABs often used data and parameters that are not readily available or easily derived from routine or event monitoring. Process-based models often needed data that described biological processes that can act as sources (photosynthesis and growth) and sinks (benthic and pelagic grazing, respiration, senescence, sedimentation, motility) on algal biomass. A range of mathematical formulations exist for representing these processes in process-based models, but those formulations rely on kinetic rate parameters that can be species-, location-, and time-specific. Unfortunately, parameter values are usually not available for the specific system, algal assemblage, and time period being modelled, so modelers are often relegated to using values from the literature or “tuning” these parameters during model calibration, a situation that can lead to “equifinality” (i.e., multiple ways for a model to obtain the “right” answer (Beven and Freer, 2001)). Some rate parameters for which a fair amount of data and/or estimation approaches are often available in the literature include maximum growth rates (Eppley, 1972), respiration rates and Chl:carbon ratios (Cloern et al., 1995; Geider, 1987), but grazing rates are frequently unavailable, particularly for benthic grazers, as observations and data for those organisms are sparse or non-existent in many systems. Moreover, invasions by groups such as dreissenid mussels in novel locations can lead to rapid changes in grazing pressure (Lucas et al., 2016), rendering site-specific grazing data from a few years ago obsolete. Two articles in our review that explored the effects of grazing rates on water quality in the Seneca River, NY, USA (Canale and Chapra, 2002; Glaser et al., 2009) and another from the River Rhine, Germany, compared nutrients and grazing pressure as relative controls of algal growth rates (Schöl et al., 2002).

Conversely, and also noted by Rousso et al. (2020) for lake systems, we found that data-driven river models tend to leverage relatively easily and routinely measured data that are proxies for complex physical and biological processes incorporated more explicitly within process-based models. For example, using turbidity measurements as a proxy for light availability or using wind direction and speed as a proxy for vertical mixing. The simplification of these processes into proxy data or rate parameters is due to the challenge of directly measuring these processes, an issue not likely to be resolved in the near term. Rousso et al. (2020) found data-driven models in lakes tended to use high-frequency *in situ* fluorescence data more than process-based models and found that increased data quantity benefits model performance for data-driven models, a pattern we also suspect is present in our review of river models.

Emerging observational technologies, such as readily available satellite data and *in situ* sensors, are filling some of the gaps in data collection and availability, providing observations of rivers at high, and previously unachievable, spatial and temporal resolution. Remote sensing of algal blooms and detection of chlorophyll is expanding, especially in recent years for CyanoHAB models in lakes (Rousso et al., 2020) and is already used to provide forecasts in several lake systems (Stauffer et al., 2019). Few of the 162 articles in this review used remote sensing imagery in their efforts, with exceptions being Son et al. (2020) who used unmanned aerial vehicles and (Pyo et al., 2021) who used hyper spectral imagery. They found that the combination of *in situ* data, multidimensional imagery, and synthetic output provided complementary effects in terms of time and spatial discrepancy in data, thereby increasing the reliability of a deep learning model. Additionally, *in situ* sensors continue to advance and provide high frequency observations for a growing number of HAB indicators, a pattern also observed for lake models (Rousso et al., 2020). Remote sensing can also expand observations to habitat types, such as side channels and backwaters, that are beyond the spatial domains typically measured using *in situ* instruments. However, discrete weekly to monthly sampling remained a common connecting thread across the 162 articles reviewed herein (Figure 5D). A final hurdle is the integration of spatial and temporal data at different frequencies and resolutions and from different sources and methods. Integrated data usage is anticipated to expand the feasibility and accuracy of models for the prediction of various water quality variables and has shown promise in marine environments (Anderson et al., 2019). Yet, expanded datasets will likely require improvements to older but still popular process-based models and data-driven techniques, in addition to a coincident demand for computing power.

#### 3.6.2 Gaps in riverine setting

This literature review has identified the most common setting of river HAB modeling over the period of our literature review: eutrophic, non-wadable rivers, likely large enough to support barge traffic, that have some sort of in-stream river obstruction, streamflow modification, or both. A similar archetype was used in a review of HAB modeling in large river systems by Xia et al. (2019), but extended here for more comprehensive understanding. Xia et al. (2019) found commonalities in river systems such as the nonlimiting role of nutrients, the highly influential role of the streamflow regime, and the potential for flow regulation to mitigate bloom formation, topics we also uncovered in our systematic literature review. However, their archetype selection of large, nutrient-rich river systems, although prevalent and represents waters that are used as drinking water sources and for recreation, only accounted for approximately a quarter of the articles in this literature review. HAB modeling is occurring across rivers with a wide variety of characteristics (e.g., Figure 3) and sizes, albeit in a limited number of countries (mainly South Korea, United States, China, and Australia) and continents (i.e., Europe and Australia). In addition to the geographic clustering of river HAB modeling around the world (Figure 2), there are two additional gaps in the current literature of river HABs modeling related to environmental setting: (1) benthic HABs and (2) stagnant side and back-channel areas.

Most articles (real and idealized settings) focused exclusively on pelagic HAB conditions (93%), reflecting the focus on non-wadable rivers. Only a small subset of 10 articles (6%) included efforts to model benthic algae. Benthic accumulations of cyanobacteria are understudied compared to pelagic blooms (Wood et al., 2020) and were not included in Rousso et al. (2020). Three studies in our review focused exclusively on benthic taxa and of those, only one (Lévesque et al., 2012) focused on cyanobacteria. Lévesque et al. (2012) modeled abundance of *Lyngbya* in flowing and impounded portions of the St. Lawrence River, Canada. The other two studies (1) modeled net primary productivity of periphytic microalgae in forested streams near Rio de Janeiro, Brazil (Neres-Lima et al., 2017) and (2) predicted *Cladophora* below a dam in Japan to improve operations (Yoshioka and Yaegashi, 2017). The remaining seven articles modeled both pelagic and benthic algal groups. One dealt specifically with cyanobacterial taxa in the benthos of urban influenced streams in Saskatchewan, Canada (Bergbusch et al., 2021). The rest were broadly focused on periphyton or benthic chlorophyll. For example, Carleton et al. (2009) modeled benthic chlorophyll *a* along with the percentage cyanobacterial biomass of sestonic algae in the shallow, eutrophic Blue Earth River in Minnesota, USA, to support the development of numeric nutrient criteria for the management of impaired river systems using the AQUATOX model.

Rivers are inherently diverse and many river HAB models overlook possible contributions from productive side or back-channel areas and the connectivity between these areas and the main channel (Giblin and Gerrish, 2020; Giblin et al., 2022). These subhabitats, and at times, upstream tributaries and structural features (e.g., dams), play an important role in creating areas with low flushing, high nutrients, and/or high residence time that influence algal accumulation (Junk et al., 1989; Reynolds and Descy, 1996), yet are poorly captured in river HAB models. We identified only 16 articles where connections to productive source areas (e.g., side pool, channel, tributary, upstream reservoir, etc.) were found to be important. Five of these articles were for idealized settings and (Grover et al., 2011; Grover et al., 2009; Hsu et al., 2013; Jäger and Borchardt, 2018; Wang, 2015; Wang et al., 2015) three of these developed theoretical models that simulated algal toxin movement from stagnant side and back-channel areas to the main channel of a large river (Grover et al., 2011; Hsu et al., 2013; Wang, 2015). This collection of articles ultimately found that a ratio of the strength of longitudinal advection to longitudinal dispersion (known as the Peclet number) was a good predictor of spatial variations in algal abundance. Lateral variation in algal abundance and toxin concentration occurred when the stagnant area was hydraulically isolated from the main channel and thus lateral exchange was weak. The rest of the articles modeled real systems, mostly using process-based approaches. For example, Yan et al. (2021) modeled transverse distribution of chlorophyll *a* and found it was most affected by a ratio of chlorophyll *a* flux between tributaries and the main channel. Higher chlorophyll concentrations were found on different sides of the channel due to influences from tributaries, changes in river width, and bends in the river. Ultimately, the authors point out few studies have explored this type of transient storage variability in rivers, which we also observed from our review, and more work needs to be done.

#### 3.6.3 Synthesis challenges

Our literature review highlights noteworthy challenges to the synthesis of historical river HAB modeling efforts, which we assert stems from (1) the diversity of models, modeling endpoints, and skill metrics used over time, and (2) poorly defined ranges of environmental characteristics due to limited descriptions of the riverine setting and the input data. Rousso et al. (2020) noted similar challenges for modeling CyanoHABs in lakes. Multiple strategic plans for HAB research articulate the need for consistent and thorough descriptions of data and models as a means for learning from previous efforts and making progress towards operational, end-user-supported forecasting and nowcasting tools (Anderson et al., 2019; Ganju et al., 2016; Stauffer et al., 2019; U.S. National Office for Harmful Algal Blooms, 2024). Furthermore, more thorough descriptions of data and models would allow model developers to identify and select optimal solutions given particular environmental settings and modeling objectives.

Across the river HAB literature, quantitative measures of the riverine environment, such as drainage area, mean annual streamflow, channel dimensions, stream order, and distance to or from an obstruction, were often absent. This is a lost opportunity because we found, despite geographic clustering within specific countries (Figure 2), the articles represent over 80 different rivers and span a range of sizes and environmental conditions. Even when quantitative information about the riverine environment is available, a variety of measures are provided that are not necessarily comparable. Inconsistent and limited reporting of location information severely impedes the development of gradients (e.g., small to large rivers, oligotrophic to hypereutrophic) and limits the ability to conduct meta-analyses and derive quantitative conclusions. For example, in our literature review we made assumptions about river size by reviewing the description of the river setting in the article and at times viewing photos of the location (or near the location) on the internet. Based on visual assessment, we coarsely assigned the river sizes as “small, wadable”, “moderate, non-wadable, likely no barge traffic”, and “large, non-wadable, possible barge traffic”. This was ultimately unsatisfying but allowed us to make some general conclusions about the size of rivers present across these articles. Rousso et al. (2020) also highlighted that lack of relevant lake-specific characteristics such as trophic status and circulation patterns in their review of lake HAB models and proposed the creation of a database containing this type of information to make it easier to compare between lakes. For rivers, at a minimum, reporting drainage area and mean annual streamflow for the river being modeled is warranted and would improve the ability to quantitatively synthesize results across a range of river sizes. Additional information like trophic status, turbidity, and hydrologic characteristics (e.g., 7-day minimum) could further improve the ability to compare models and results across river systems.

No single “standard” modeling approach was identified during our review. We found a diversity of model structures (Table 4 and Table SM-4), input variables (Figure 4 and Figure 5), model complexity (e.g., process-based models, Figure 7), model endpoints, and model validation approaches (Figure 8). For example, model skill was challenging to compare across articles because, in many cases, a quantitative skill assessment was either not performed or only minimally explained, a finding that is apparent across lake models as well (Rousso et al., 2020). It was also sometimes unclear which output variables (e.g., the HAB-related modeling endpoint or an additional output like nutrient concentrations) were used to calculate the skill metric if one was reported. Moreover, although a few skill metrics were somewhat common (e.g., R^2^, RMSE), a vast array of metrics was employed across the reviewed articles, including deviance, Willmott’s Score, mean error, mean absolute error, accuracy, specificity, posterior predictive p-value, misclassification rate, Nash-Sutcliffe efficiency, among others. Finally, two papers could present a particular metric (e.g., R^2^) for a specific output variable (e.g., chlorophyll *a* concentration), but those two metrics may not be entirely comparable because they may have been computed relative to different kinds and numbers of observational data (e.g., timeseries at a single location, a single time across multiple locations) or for different kinds of model runs (e.g., a calibration run for which model coefficients were adjusted to optimize model-observation match versus a “validation” run for which coefficients were already independently determined). Ultimately, the lack of consistent model skill metrics hinders model comparison and the ability to identify promising approaches.

Modeling decisions, including the choice of endpoints (and input variables), were often shaped by data availability and practical considerations, which in turn influence the interpretation and application of model results. Gorney et al. (2023) and Ho and Michalak (2015) describe the nuance of defining a HAB and how a selected HAB definition (whether explicit or implicit) determines the hypothesis that can be tested and how model results may be used. Many of the 162 articles in our review used chlorophyll over more specific, taxa-related endpoints, likely because of the comparative ease of the collection and laboratory analysis of this measure. Furthermore, few articles used thresholds or modeling endpoints that were explicitly related to a perceived, potential, or actual harm, such as algal toxin concentration or exceedance of algal biomass above a recreational threshold. This topic was not discussed for lake models in Rousso et al. (2020), where the discussion of modeling endpoints focused exclusively on chlorophyll *a,* cell concentration, biovolume, and biomass. The wide range of endpoints used across the 162 river-focused articles underscores the difficulty of synthesizing model output and gauging a model’s utility; a more uniform approach to reporting this information is warranted.

Relatedly, some of the decisions that led to the use of certain input variables were also likely based on data availability and accessibility which ultimately influences which predictors were deemed important upon the completion of modeling. Potentially important processes were only rarely included (as an equation or a proxy variable) in models. For instance, processes such as migration of cells, grazing, toxin production, benthic-pelagic interactions, or exchange between side or back-channel areas with a mainstem were often neglected. Measuring these processes in real systems is a challenge that results in a lack of observational data. There is uncertainty about how to represent some of these processes mathematically. Yet, identifying the importance of a process or variable depends on its inclusion in the model. Some of the differences between the input variables (Figure 4), simulated processes in process-based models (Figure 7), and important predictors identified by the authors (Figure 9) demonstrate this issue. For example, “algal processes” was used as an input variable in just 23% of the data-driven modeling articles (Figure 4) and was found to be important in the same percentage of data-driven modeling articles (23%, Figure 9). In contrast, process-based models include algal processes as input data or an internal mathematical process in over 80% of the process-based modeling articles (Figure 7) and algal processes was found to be the most important variable across process-based modeling articles (38% of all process-based modeling articles; Figure 9). The discrepancy in the importance of algal processes for modeling river HABs between data-driven versus process-based model approaches likely stems from whether the data (or processes) are included in the models in the first place. Data used during model development and as input variables often reflects the difficulty associated with measuring, analyzing, and incorporating that observational information into a model, and these decisions ultimately dictate the insights that can be gained from the model results. This begs the question: Are we using the best modeling inputs and endpoints or just the easiest ones?

### 3.7 Opportunities for advancing river HAB modeling

Although challenges abound, our review also highlights opportunities for advancing river HAB modeling. In particular, there is much to learn from estuarine HAB models as these model formulations include riverine processes and have been advanced by many researchers over recent decades. Additionally, forecasting river HABs is a promising way forward that can leverage theoretical or system-specific models and apply these towards user-relevant management goals. Finally, community-developed common datasets may serve to encourage rapid development of river HAB models that can be readily evaluated and compared.

#### 3.7.1 Learning from estuarine HAB models

It is crucial to improve our understanding of HAB drivers in freshwater and coastal systems and to view HABs in these domains as connected, both mechanistically and with respect to symbiotic approaches for studying them. Estuaries—aquatic systems where fresh river water meets salty ocean water—are a physical representation of that connection, and cyanotoxins produced in upstream source waters can propagate to estuaries and be taken up by mollusks, fish, and mammals and thus lead to human health risks (Miller et al., 2010; Preece et al., 2017). In our review, 21 articles indicated an estuarine context (Table SM-1). Most used process-based modeling (76% of the estuarine modeling articles) and around half of these articles modeled a system in the United States (52%). Although estuarine HAB modeling represented a small portion of articles, they provide an interesting view into what advanced and specific models can offer to the river HAB modeling community.

Estuaries can, in some cases, function like rivers and, for that reason, many of the findings and modeling tools from studies of estuarine phytoplankton dynamics could be extended and applied to rivers. For example, if river inflows to an estuary are high enough to dominate over tidal influences in governing net horizontal transport, the estuary’s hydrodynamics may, in the tidally averaged sense, approximate that of a river, particularly if the estuary is relatively narrow, such as a tidal river. Many of the simple (analytical) estuarine phytoplankton models (e.g., Qin and Shen, 2021; Wang et al., 2019c) could be applied in rivers; in fact, because of their simplifying assumptions regarding geometry and hydrodynamics, some of those models may indeed be more appropriate for rivers than for estuaries (e.g., Lucas and Thompson, 2012; Lucas et al., 2009b). Analogous to the thermal stratification found in some rivers or run-of-the-river reservoirs, estuaries typically experience some degree of salinity stratification (Fischer et al., 1979) that may be enhanced by thermal stratification (Vroom et al., 2017). The models and approaches developed to explore the influence of stratification and vertical turbulent mixing on estuarine algal blooms (e.g., Burchard et al., 1999; Koseff et al., 1993; Lucas et al., 1998) could be applied to investigate physical-biological bloom controls operating in the vertical dimension of rivers (e.g., interactions between stratification, turbulent mixing, cell sedimentation, migration, and buoyancy).

Another relevant theme is the lateral transport of algal biomass from productive side source areas (shoals, in the case of drowned river estuaries) to the deeper main channel (Engel et al., 2025; Lucas et al., 2016; Lucas et al., 2009a; May et al., 2003); such modeling tools developed for studying the estuarine realm could be adapted for the study of algal biomass exchange between potentially productive riverine backwaters or side storage areas and the mainstem. Finally, much of the multi-dimensional hydrodynamic-ecological software that has been applied to characterize phytoplankton growth, loss, and two- or three-dimensional transport in estuaries is general enough to capture relevant processes in rivers (as noted in Table 4) and other water body types. Examples within this review include EFDC (Environmental Fluid Dynamics Code; Qin and Shen, 2019) and MIKE (Lubello et al., 2025). Other models such as Delft3D (Castro-Olivares et al., 2024), Delft3D-FM (Flexible Mesh; White et al., 2021), and SCHISM/CoSiNE (Wang et al., 2020) also have this capability.

#### 3.7.2 Forecasting and Operationalization

There are fewer examples of forecasting HABs for inland waters compared to marine systems, and even less in river than lake systems (U.S. National Office for Harmful Algal Blooms, 2024). There are exceedingly few sustained operational forecasting models for HABs in lotic systems, though notable exceptions include the work of Park et al. (2021) in the Changnyung-Haman Reservoir, South Korea and Nietch et al. (2022) for the entire 1579 km of the Ohio River (United States). Our literature review identified 21 articles that developed forecasting models. Not all of these were operational; many were used for research purposes in the past (i.e., how well did forecasts perform historically). Notably, forecasting efforts were heavily concentrated in South Korea, which accounted for 14 of the 21 articles, whereas the second ranked country in forecasting applications, Australia, contributed only three articles. Among all forecasting applications, predicting HABs seven days into the future was the most common lead time. The shortest lead time observed was one day, whereas the longest extended to 55 days Nietch et al. (2022). There is a growing consensus that regional modeling efforts, that can be adapted to specific systems, is likely a productive path forward for inland water (U.S. National Office for Harmful Algal Blooms, 2024), and perhaps especially for rivers.

Despite these efforts, there is currently a lack of operationalization in forecasting models. Few studies have moved toward implementing early warning systems or actionable measures, such as information to support beach closures or drinking water treatment decisions. We suspect that the limited application of forecasting is due to the challenges associated with operationalizing models for routine predictions. Unlike weather forecasting, there is currently no standardized cyberinfrastructure for river HABs (or other environmental forecasting for that matter) that researchers can use. Furthermore, transitioning models to operation relies on continued investments in research, monitoring, and cyberinfrastructure (U.S. National Office for Harmful Algal Blooms, 2024). As a result, researchers and organizations develop their own separate forecasting systems. This is also reflected in the diversity of models and modeling approaches found in our review, which is also apparent for lake models (Rousso et al., 2020). This fragmentation or disconnection often means that river HABs forecasts are produced manually or using highly site-specific, independently developed methods, which hampers a collective ability to efficiently generate important ecological predictions and deepen understanding of the environment.

Nonetheless, there are promising avenues for addressing these issues, including examples of national-scale monitoring networks and reporting systems, in addition to community-developed tools. South Korea uses an extensive national-scale database of algal information and provides real-time HAB risk categories based on observed cyanobacteria cell density for multiple locations on all major South Korean rivers (http://water.nier.go.kr/web/algaeStat?pMENU_NO=195). This system is a notable example of operational river HAB nowcasting and provides quantitative data along with approachable indicators of risk for water supply and recreation. In terms of forecasting, operational community-developed tools could facilitate increased forecasting of river HABs in the future. Synergistic partnerships between research groups and with funding agencies provide a means for sharing resources (U.S. National Office for Harmful Algal Blooms, 2024). For instance, the Ecological Forecasting Initiative has established shared cyberinfrastructure to support forecasting challenges (Thomas et al., 2023), including a river chlorophyll forecasting challenge co-hosted by the Ecological Forecasting Initiative and the U.S. Geological Survey (https://waterdata.usgs.gov/blog/habs-forecast-challenge-2024/). We believe that community-driven cyberinfrastructure will promote the iterative improvement of river HABs forecasts, ultimately supporting decision-making and effective water management.

#### 3.7.3 Towards Common Datasets

Most articles in our literature review used unique datasets curated for a specific study, either based on location, application, or both. This variability poses challenges for collectively advancing river HABs modeling as modeling endpoints, goals, and available input variables differ across datasets (Figures 4, 5, SM-1, and SM-2). Consequently, applying various types of models to the same dataset becomes difficult. When one model outperforms another in different analyses, it is often unclear whether the observed differences are because of genuine variations in model skill, the specific modeling task, or the dataset used in the modeling process. Rousso et al. (2010) noted that there were few studies that applied multiple modeling approaches to the same lake. We identified a similar pattern in our review, with the exception of the Nakdong River in South Korea. As a focal point for many river HAB models, there is potential for additional synthesis of these articles and modeling approaches. Nevertheless, standardizing datasets for model training, calibration, and evaluation could accelerate progress in environmental modeling, including for river HABs. Based on the success of South Korea’s Water Environment Information System (http://water.nier.go.kr/web) and the many river HAB modeling articles leveraging these data, it would be prudent for a standardized dataset to include cyanobacteria cell counts for multiple potential cyanotoxin-producing taxa, ideally at a weekly frequency, along with chlorophyll concentrations and potentially important predictor variables. Based on Figure 9, additional data could include N and P concentrations (in various forms), measures of light availability, water temperature, and streamflow (or velocity). Supporting information capturing algal processes such as predation, cell motility, resting states, among others, would also be beneficial.

In machine learning, standardized datasets are referred to as “benchmark” datasets, and there are increasingly more examples of these used for environmental applications. For instance, streamflow prediction models have successfully used global benchmark datasets (Kratzert et al., 2023), allowing researchers to (1) focus on model improvement rather than collecting and/or curating datasets, and (2) compare model performance to other models applied to the same data. Additionally, a freshwater forecasting challenge identified that process-based models outperformed data-driven models for forecasting water temperature, whereas the opposite was true for forecasting dissolved oxygen (Olsson et al., 2025). Perhaps this can be interpreted as process-based models excelling at deterministic, physics-based processes whereas data-driven models better capture biological process that are more influenced by stochastic processes. Recent efforts have been made to curate river chlorophyll and ancillary datasets at national scales (Fernandez et al., 2025; Spaulding et al., 2024), which could substantially enhance riverine chlorophyll modeling in support of more comprehensive HAB studies. Given the global prevalence of river HABs highlighted in our review, we are hopeful that a comprehensive global benchmark dataset will soon be developed, similar to those already available for streamflow (Kratzert et al., 2023).

## 4. Conclusions

Our review encompassed 162 articles spanning nearly five decades, modeling over 80 rivers globally, and identified several challenges and opportunities for river HAB modeling. Three such challenges are (1) the limited availability and integration of key and emerging data sources, (2) geographic and ecological biases in modeling efforts, and (3) difficulties in comparing and synthesizing results due to inconsistent reporting of river characteristics (e.g., drainage area, mean annual streamflow, etc.) and model evaluation metrics. We also note few modeling articles addressed algal toxins, connections to side channels and back waters, or benthic algae. Despite these existing challenges and gaps, promising progress of river HAB modeling is emerging through estuarine modeling efforts, South Korea’s nowcasting algal alert system, and community-developed tools spearheaded by groups like the Ecological Forecasting Initiative.

Ongoing changes in nutrient and climatic conditions underscore the benefits of a broader modeling focus and enhanced, or at least sustained, monitoring. Although nutrients, elevated water temperatures, and low-flow conditions are well-established drivers of HABs in rivers (Griffith and Gobler, 2020; Schmadel et al., 2024), changing environmental conditions have led to increased HAB occurrence in unexpected environments, including benthic habitats (Wood et al., 2020), coastal estuaries (Preece et al., 2017), and low nutrient systems (Reinl et al., 2021). There is a need and an opportunity to model a wider range of systems with emerging algal issues, especially where causal factors are less well understood. This includes geographic areas where river HAB modeling is largely absent, such as the southern hemisphere and equatorial regions. Furthermore, despite many models being used to test scenarios, few studies emphasized management, provided strong recommendations regarding data collection, or developed strong foundations for operationalized forecasts. Moving forward, the development of integrated approaches, enhancement of model applicability across diverse environments, and strengthening of the connection between research findings and practical management strategies could improve outcomes in addressing HABs in riverine systems and beyond.

## Supporting information

Supporting Materials-2: Comprehensive Bibliography

## Acknowledgements

We thank Maria (Masha) Marionkova for completing initial pulls from scientific databases and initial work on the data collection form; we thank Gabriella Zuccolotto and Sarah Stackpoole for providing critical reviews of a few articles. This work was completed as part of the U.S. Geological Survey (USGS) Proxies Project, an effort supported by the Water Mission Area (WMA) Water Quality Processes program to develop estimation methods for PFAS, harmful algal blooms, and metals, at multiple spatial and temporal scales. Additional support for this work was from the USGS Integrated Water Availability Assessments (IWAAs) Program, which examines the spatial and temporal distribution of water quantity and quality in both surface and groundwater, as related to human and ecosystem needs and as affected by human and natural influences. Any use of trade, firm, or product names is for descriptive purposes only and does not imply endorsement by the U.S. Government.

## Data Release

Data gathered from literature and presented in this manuscript are available at: Gorney, R.M., Zwart, J.A., Lucas, LV., and Murphy, J.C., 2025, Data from a systematic literature review of forecasting and predictive models for harmful algal blooms in flowing waters: U.S. Geological Survey data release, https://doi.org/10.5066/P1JWCCXF.

