## Supporting Materials-1: Additional Text, Tables, and Figures for "A systematic literature review of forecasting and predictive models of harmful algal blooms in flowing waters"

This information product has been peer reviewed and approved for publication as a preprint by the U.S. Geological Survey.

### Extended Model Type Descriptions

#### Process-based models

Process-based (i.e., mechanistic) models include mathematical representations of some or many of the individual processes mechanistically linking output or response variables (effects) to input or predictor variables (causes) in explicit mass balances (Lucas et al., 2025). There is a range of ways in which an individual process—such as grazing of algae by zooplankton—may affect mass and be included in a model. For example, a simple first-order loss rate may be specified by the user and included in the governing mass-balance equation for algal biomass (e.g., Engel et al., 2025; Lucas et al., 1999). Such mathematical relations may be based on first principles (e.g., conservation of mass or momentum), empirical relations (e.g., photosynthesis-irradiance relations for calculating algal production and growth), or both. Process-based models span a broad range of complexity, from numerical models that solve systems of multi-dimensional partial differential equations describing realistic conditions, to analytical models that offer exact solutions. Numerical models implement “numerical methods”, which are approaches for estimating solutions to mathematical problems when an exact solution is not possible. A relevant river example would be for the solution of the time-dependent advection-dispersion-reaction equation for algal biomass, *B*, under realistic spatially and temporally varying conditions:

$$\frac{\partial B}{\partial t} + u\frac{\partial B}{\partial x} + v\frac{\partial B}{\partial y} + w\frac{\partial B}{\partial z} = K_x\frac{\partial^2 B}{\partial x^2} + K_y\frac{\partial^2 B}{\partial y^2} + K_z\frac{\partial^2 B}{\partial z^2} + \mu_{growth}B - \mu_{loss}B$$

where *x*, *y*, and *z* are the longitudinal, lateral, and vertical dimensions; *u*, *v*, and *w*, respectively, are current velocities in those three directions; $K_x$, $K_y$, and $K_z$, respectively, are mixing or dispersion coefficients in the *x*, *y*, and *z* directions; $\mu_{growth}$ is a first-order growth rate; $\mu_{loss}$ is a first-order loss rate, which may include grazing, death, and/or other loss processes (Lucas, 2010). In a realistic aquatic system, velocities, mixing coefficients, and growth and loss rates will all vary spatially and temporally, rendering the above equation impossible to solve exactly for *B*. However, numerical methods such as finite difference, finite volume, or finite element schemes are available to provide approximate solutions. Such methods “discretize” time and space, respectively, into time steps and a grid of points, cells, volumes, or elements (e.g., Qin and Shen, 2019).

Analytical models, are those which solve mathematical problems with an exact (or "closed form") solution, rendering numerical techniques unnecessary. Usually, analytical models describe simplified or idealized situations as necessary to solve the mass balance. For example, many simplifying assumptions dropped terms from the above differential equation (e.g., one-dimensional, steady-state, no dispersion), and assumes that velocity, growth and loss are spatially uniform, the analytical solution may become simple and be evaluated with the following direct algebraic expression for *B*:

$$B(x) = B_{in} exp\left(\frac{\mu_{growth} - \mu_{loss}}{u} x\right)$$

where $B_{in}$ is the algal biomass concentration entering the system at the upstream end, and *B*(*x*) is the concentration at longitudinal location *x* (Lucas et al., 2009). As long as one knows $B_{in}$, $\mu_{growth}$, $\mu_{loss}$, and *u*, this simple equation can be easily solved for *B*(*x*).

### Data-driven models

Data-driven (i.e., statistical, empirical, machine learning, or deep learning) models are generally used to predict continuous or categorical outcomes based on a set of features or predictor variables (Lucas et al., 2025). Such models do not predefine system processes or mechanisms; instead, they allow the data to reveal relations between input and output variables. Data-driven models range from simple regression methods to more complex machine learning approaches, such as random forests and neural networks (Lucas et al., 2025).

Simple data-driven models, herein referred to as a "simple statistical" subtype (e.g., linear regression), offer practical advantages in terms of data requirements compared to process-based or more complex data-driven models. These models do not require estimates of process rates or constants, enabling straightforward regression of an output variable against an input variable using relatively few data points. Bayesian methods are a step up from simple statistical methods and can incorporate prior information and established relations, which enhances predictive performance, particularly in data-sparse systems. Furthermore, Bayesian methods excel at estimating predictive uncertainty and facilitating adaptive learning by easily incorporating new data as it becomes available.

More complex data-driven techniques, referred to as "machine learning" data-driven subtype (e.g., random forest, neural networks), can manage multiple correlated variables and capture non-linear relations among input and output variables. Additionally, variable importance analyses can help modelers assess the relative contributions and influences of input variables. Neural networks represent the most flexible yet data-intensive option for data-driven modeling. They offer significant advantages over tree-based methods, particularly in their ability to incorporate diverse data types and capture complex spatial and temporal relations within a system. For instance, convolutional neural networks and recurrent neural networks can be combined to analyze the spatial and temporal dynamics of algal blooms in inland waters, using satellite imagery and weather data alongside *in situ* sampling. However, these complex data-driven models often require large datasets, sometimes thousands of observations, to accurately learn intricate patterns, which can pose challenges for environmental monitoring efforts, particularly when capturing low-frequency events, such as HABs in rivers.

Table SM-1. Reference lists for articles comprising topical collections of interest not easily parsed from the complete dataset (Gorney et al., 2025). Full bibliographic citations are available in Supporting Materials-2: Bibliography.

| Topic | Article citations |
| --- | --- |
| Validation papers (n = 24) | (Ahn et al., 2021; Grover et al., 2011; Grover et al., 2017; He et al., 2021b; Kim et al., 2021a; Kim et al., 2022a; Kim et al., 2022b; Kim et al., 2018; Kim et al., 2019; Kim et al., 2020; Kim et al., 2021b; Kim et al., 2022c; Lee and Lee, 2018; Lévesque et al., 2012; Liao et al., 2021; Maier et al., 1998; Nietch et al., 2022; Pathak et al., 2021; Pyo et al., 2020; Shan et al., 2022; Su et al., 2022; Teles et al., 2006; Wang et al., 2019; Yi et al., 2018) |
| Peer reviewed government reports (n = 5) | (Rounds and Wood, 2001; Rounds et al., 1999; Smith et al., 2022; Sullivan and Rounds, 2018; Sullivan et al., 2011) |
| Modeling endpoint: Algal toxins (n = 8) | (Abbas, 2015; Grover et al., 2011; He et al., 2021a; He et al., 2021b; Hsu et al., 2013; Shan et al., 2022; Wang, 2015; Wang et al., 2015) |
| Used high and low frequency data in modeling effort (n = 12) | (Bormans and Condie, 1998; Crossman et al., 2021; Harvey et al., 2024; Kim et al., 2021a; Kim et al., 2023; Loewenthal et al., 2012; Pathak et al., 2021; Savoy and Harvey, 2023; Schöl et al., 2014; Stringfellow et al., 2009; Wang et al., 2019; Waylett et al., 2013) |
| Estuarine articles (n = 21) | (Arhonditsis et al., 2007; Cerco et al., 2004; Crossman et al., 2021; Ducharne, 2008; Jeong et al., 2003; Kim et al., 2017; Kim et al., 2022b; Lee et al., 2021; Liu et al., 2000; Lung, 1988; Lung, 1986; Pinckney et al., 1997; Savoy and Harvey, 2023; Scharfe et al., 2009; Schöl et al., 2014; Smith et al., 2022; Wan et al., 2022; Wang and Zhang, 2020; Wang et al., 2020; Wang et al., 2019) |

Table SM-2. Copy of inclusion/exclusion criteria used during Phase 2 screening process. Only included peer reviewed publications (no proceedings, abstracts, conference papers, etc.). Screening was completed sequentially at three different levels: (a) title and keywords, (b) abstract, and (c) full text, according to three criteria described below. Articles needed to meet all 3 of the inclusion criteria and contain none of the exclusion criteria to be included in the final set of articles.

| Inclusion criteria | Exclusion criteria |
| --- | --- |
| 1) The research must have been conducted in freshwater flowing environments such as rivers, streams, creeks, canals, channels, and ditches whether natural, constructed, modified, or managed. May include run-of-river reservoirs, lock and dam pools, and tidally influenced river environments. Instream reservoirs and rivers in estuarine settings | • Excludes research exclusively focused on lakes, reservoirs, marine environments, and estuaries.<br>• Excludes ambiguous riverine settings (e.g., tidal, estuarine, in-stream reservoirs, off channel waters, wetlands) where the corresponding model includes processes that are clearly estuarine (e.g., salinity stratification, gravitational circulation, or |

| Inclusion criteria | Exclusion criteria |
| --- | --- |
| are also included if the corresponding model contains clearly riverine processes (e.g., advective flow). Models of idealized settings are also included. | tidally driven vertical mixing) or lacks clearly riverine processes.<br>• Excludes research where rivers are included only as inputs to the model and not a focus of the modeling effort. |
| 2) Models must have been developed or implemented for prediction (i.e., between observations over time or across space), hindcasting, nowcasting, forecasting, or scenario or sensitivity analyses (e.g., related to climate change or management decisions). | • Excludes models that do not produce predictions, hindcasts, forecasts, or scenarios (e.g., models where only the fitted coefficients are used for interpretation, and predictions are not made at novel locations or periods of time).<br>• Excludes studies with purely conceptual models (i.e., models lack any mathematical or statistical components).<br>• Excludes models or algorithms that only use remotely sensed spectra or spectral indices to estimate chlorophyll or other algal endpoints. Excludes models or algorithms developed for the sole purpose of validating a spectral index. Does not exclude models that incorporate remotely sensed information along with other observations to predict blooms or algal dynamics. |
| 3) Modeling endpoints must be measures of algal composition, abundance, biomass, presence, or toxicity (e.g., community composition, cell counts, biovolume, toxin concentrations) or related proxies (e.g., chlorophyll, phycocyanin). This includes models that predict phytoplankton primary production and algal cellular nutrient content. Novel proxies will also be considered (e.g., primary productivity or oxygen dynamics) as long as the stated purpose is for algal bloom prediction. Models providing categorical output or estimates of bloom risk or probability are also included. | • Excludes research related to specific harms which do not include algal biomass prediction or forecasting within the modeling framework. For example, excludes studies focused exclusively on changes to the food web (i.e., the effect of changes in algae composition or biomass on other organisms), algal toxin health impacts on humans or animals, and effects of a bloom occurrence on the local economy.<br>• Excludes studies that only evaluate environmental conditions that could affect algal dynamics (e.g., only nutrients, sediment, or temperature). |

Table SM-3. Number (and percent) of system-specific articles by country (n = 151) and number and percent by model type (system-specific data-driven n = 60, process-based n = 84, other n = 7)

| Country | All articles, n | All articles, percent | Data-driven articles, n (% of data-driven articles) | Process-based articles, n (% of process-based articles) | Other model articles, n (% of other model articles) |
|---|---|---|---|---|---|
| South Korea | 40 | 26 | 26 (43%) | 14 (17%) | --- |
| United States | 32 | 21 | 9 (15%) | 22 (26%) | 1 (14%) |
| China | 18 | 12 | 8 (13%) | 8 (9.5%) | 2 (29%) |
| Australia | 10 | 6.6 | 5 (8.3%) | 3 (3.6%) | 2 (29%) |
| France | 9 | 6 | --- | 9 (11%) | --- |
| Germany | 8 | 5.3 | 1 (1.7%) | 7 (8.3%) | --- |
| United Kingdom | 7 | 4.6 | --- | 6 (7.1%) | 1 (14%) |
| Canada | 6 | 4 | 4 (6.7%) | 2 (2.4%) | --- |
| Belgium | 4 | 2.6 | --- | 4 (4.8%) | --- |
| Portugal | 2 | 1.3 | 2 (3.3%) | --- | --- |
| Brazil | 2 | 1.3 | 2 (3.3%) | --- | --- |
| Russia | 2 | 1.3 | 2 (3.3%) | --- | --- |
| Hungary | 2 | 1.3 | 1 (1.7%) | 1 (1.2%) | --- |
| South Africa | 2 | 1.3 | --- | 2 (2.4%) | --- |
| Finland | 1 | 0.66 | --- | --- | 1 (14%) |
| Poland | 1 | 0.66 | --- | 1 (1.2%) | --- |
| Romania | 1 | 0.66 | --- | 1 (1.2%) | --- |
| Ukraine | 1 | 0.66 | --- | 1 (1.2%) | --- |
| Japan | 1 | 0.66 | --- | 1 (1.2%) | --- |
| Spain | 1 | 0.66 | --- | 1 (1.2%) | --- |
| Not reported | 1 | 0.66 | --- | 1 (1.2%) | --- |

Table SM-4. Number of data-driven articles (n=60) across 43 distinct data-driven techniques.

| Data-driven modeling techniques | Data-driven articles, n | Data-driven articles, percent |
|---|---|---|
| artificial neural network | 11 | 18 |
| multiple linear regression | 7 | 12 |
| other | 5 | 8.3 |
| random forest | 4 | 6.7 |
| convolutional neural network | 3 | 5 |
| long short-term memory | 3 | 5 |
| regression tree | 3 | 5 |
| AdaBoost | 2 | 3.3 |
| Bayesian structural equation model | 2 | 3.3 |
| Markov chain | 2 | 3.3 |
| Not reported | 2 | 3.3 |
| XGBoost | 2 | 3.3 |
| bootstrap aggregation | 2 | 3.3 |
| classification tree | 2 | 3.3 |
| decision tree | 2 | 3.3 |
| genetic programming | 2 | 3.3 |
| kernel density estimation | 2 | 3.3 |
| logistic regression | 2 | 3.3 |
| radial basis function network | 2 | 3.3 |
| recurrent neural network | 2 | 3.3 |
| Bayesian hierarchical model | 1 | 1.7 |
| Bayesian inverse model | 1 | 1.7 |
| Bayesian model averaging | 1 | 1.7 |
| Chi-square automatic interaction detection | 1 | 1.7 |
| Getis-Ord Gi* | 1 | 1.7 |
| K-means clustering | 1 | 1.7 |
| adaptive neuro-fuzzy inference system | 1 | 1.7 |
| autoregressive integrated moving average | 1 | 1.7 |
| deep belief network | 1 | 1.7 |
| evolutionary algorithm | 1 | 1.7 |
| extreme learning machine | 1 | 1.7 |
| gated recurrent unit | 1 | 1.7 |
| generalized additive model | 1 | 1.7 |
| generalized linear model | 1 | 1.7 |
| generalized regression neural network | 1 | 1.7 |
| genetic algorithm | 1 | 1.7 |
| geographically weighted regression | 1 | 1.7 |

| Data-driven modeling techniques | Data-driven articles, n | Data-driven articles, percent |
|---|---|---|
| hidden Markov model | 1 | 1.7 |
| partial least squares path model | 1 | 1.7 |
| principal component analysis | 1 | 1.7 |
| reverse time attention | 1 | 1.7 |
| self-organizing map | 1 | 1.7 |
| support vector machine | 1 | 1.7 |

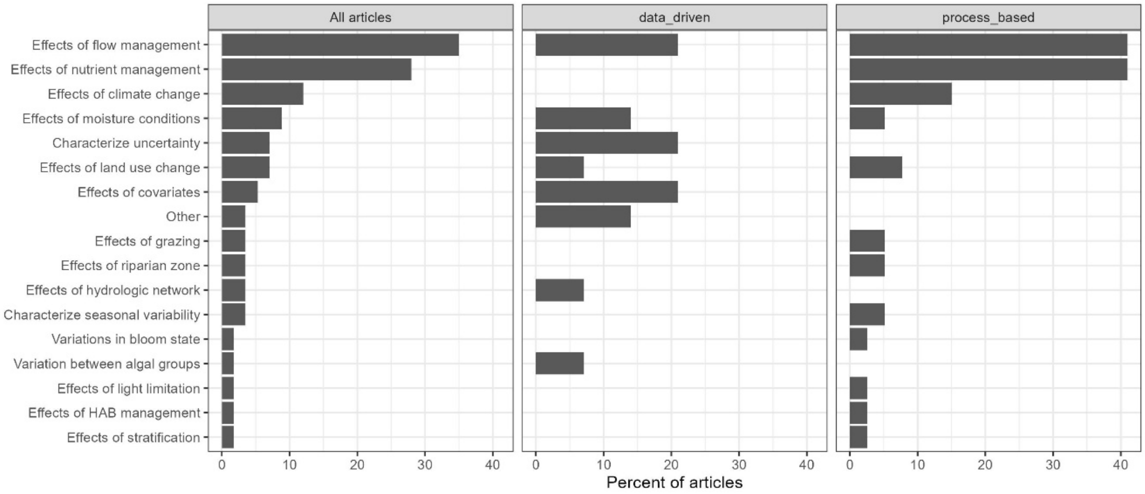


Figure SM-1. Number and percent of articles that explored a given scenario, for any articles that used a model to test a scenario (n = 57), only data-driven articles (n = 14), and only process-based articles (n = 39). [HAB, harmful algal bloom]

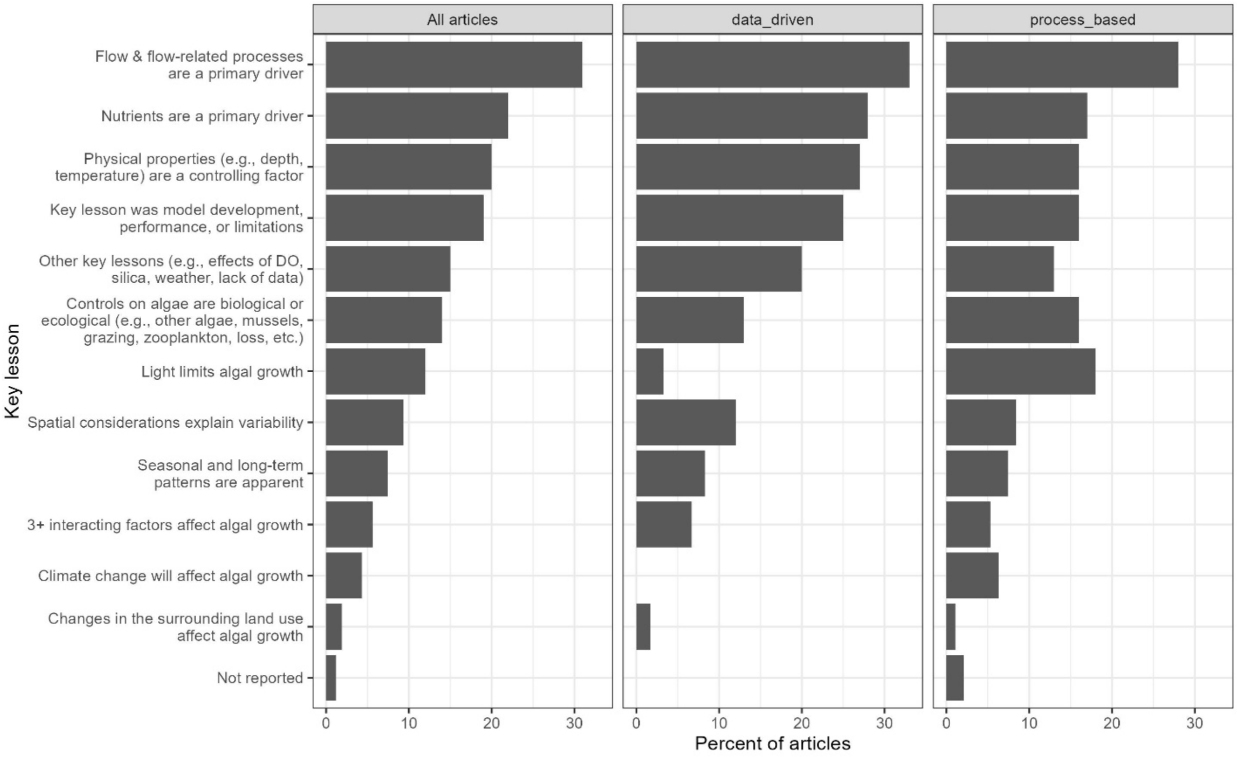


Figure SM-2. Key lessons identified by authors, as a percent of all articles (n = 162), data-driven modeling articles (n = 60), and process-based modeling articles (n = 96). Note, the “All articles” panel includes 7 articles with models classified as “other”. [DO, dissolved oxygen]
