## Supporting Materials-2: Comprehensive Bibliography for "A systematic literature review of forecasting and predictive models of harmful algal blooms in flowing waters"

This information product has been peer reviewed and approved for publication as a preprint by the U.S. Geological Survey.

Abbas, S. 2015. Dynamical analysis of a model of harmful algae in flowing habitats with variable rates. Nonlinear Analysis: Real World Applications 22, 16-33. 10.1016/j.nonrwa.2014.06.001.

Ahn, J.M., Kim, B., Jong, J., Nam, G., Park, L.J., Park, S., Kang, T., Lee, J.K. and Kim, J. 2021a. Predicting Cyanobacterial Blooms Using Hyperspectral Images in a Regulated River. Sensors 21(2), 530. 10.3390/s21020530.

Ahn, J.M., Kim, J., Park, L.J., Jeon, J., Jong, J., Min, J.-H. and Kang, T. 2021b. Predicting Cyanobacterial Harmful Algal Blooms (CyanoHABs) in a Regulated River Using a Revised EFDC Model. Water 13(4). 10.3390/w13040439.

Arhonditsis, G.B., Paerl, H.W., Valdes-Weaver, L.M., Stow, C.A., Steinberg, L.J. and Reckhow, K.H. 2007a. Application of Bayesian structural equation modeling for examining phytoplankton dynamics in the Neuse River Estuary (North Carolina, USA). Estuarine, Coastal and Shelf Science 72(1-2), 63-80. 10.1016/j.ecss.2006.09.022.

Arhonditsis, G.B., Stow, C.A., Paerl, H.W., Valdes-Weaver, L.M., Steinberg, L.J. and Reckhow, K.H. 2007b. Delineation of the role of nutrient dynamics and hydrologic forcing on phytoplankton patterns along a freshwater–marine continuum. Ecological Modelling 208(2-4), 230-246. 10.1016/j.ecolmodel.2007.06.010.

Bergbusch, N.T., Hayes, N.M., Simpson, G.L. and Leavitt, P.R. 2021. Unexpected shift from phytoplankton to periphyton in eutrophic streams due to wastewater influx. Limnology and Oceanography 66(7), 2745-2761. 10.1002/lno.11786.

Billen, G., Garnier, J. and Hanset, P. 1994. Modelling phytoplankton development in whole drainage networks: the RIVERSTRAHLER Model applied to the Seine river system. Hydrobiologia 289(1-3), 119-137. 10.1007/bf00007414.

Bormans, M. and Condie, S.A. 1998. Modelling the distribution of *Anabaena* and *Melosira* in a stratified river weir pool. Hydrobiologia 364(1), 3-13. 10.1023/a:1003103706305.

Bruns, N.E., Heffernan, J.B., Ross, M.R.V. and Doyle, M. 2022. A simple metric for predicting the timing of river phytoplankton blooms. Ecosphere 13(12). 10.1002/ecs2.4348.

Bucci, V., Nunez-Milland, D., Twining, B.S. and Hellweger, F.L. 2011. Microscale patchiness leads to large and important intraspecific internal nutrient heterogeneity in phytoplankton. Aquatic Ecology 46(1), 101-118. 10.1007/s10452-011-9384-6.

Bussi, G., Whitehead, P.G., Bowes, M.J., Read, D.S., Prudhomme, C. and Dadson, S.J. 2016. Impacts of climate change, land-use change and phosphorus reduction on phytoplankton in the River Thames (UK). Science of The Total Environment 572, 1507-1519. 10.1016/j.scitotenv.2016.02.109.

Cabecinha, E., Lourenço, M., Moura, J.P., Pardal, M.Â. and Cabral, J.A. 2009. A multi-scale approach to modelling spatial and dynamic ecological patterns for reservoir's water quality management. Ecological Modelling 220(19), 2559-2569. 10.1016/j.ecolmodel.2009.06.011.

Canale, R.P. and Chapra, S.C. 2002. Modeling Zebra Mussel Impacts on Water Quality of Seneca River, New York. Journal of Environmental Engineering 128(12), 1158-1168. 10.1061/(asce)0733-9372(2002)128:12(1158).

Carleton, J.N., Park, R.A. and Clough, J.S. 2009. Ecosystem modeling applied to nutrient criteria development in rivers. Environmental Management 44(3), 485-492. 10.1007/s00267-009-9344-2.

Cerco, C.F., Noel, M.R. and Tillman, D.H. 2004. A practical application of Droop nutrient kinetics (WR 1883). Water Research 38(20), 4446-4454. 10.1016/j.watres.2004.08.027.

Cho, Y. and Kim, Y. 2019. Extracting the factors influencing chlorophyll-a concentrations in the Nakdong River using a decision tree algorithm. Desalination and Water Treatment 157, 195-208. 10.5004/dwt.2019.24195.

Chung, S.-W., Chong, S.-A. and Park, H.-S. 2016. Development and Applications of a Predictive Model for Geosmin in North Han River, Korea. Procedia Engineering 154, 521-528. 10.1016/j.proeng.2016.07.547.

Chung, S.W., Lee, H. and Jung, Y. 2008. The effect of hydrodynamic flow regimes on the algal bloom in a monomictic reservoir. Water Science and Technology 58(6), 1291-1298. 10.2166/wst.2008.482.

Claudson, R.M. 1975. The digital simulation of river plankton population dynamics. Communications of the ACM 18(9), 517-523. 10.1145/361002.361012.

Cloot, A. and Roux, G.L. 1997. Modelling algal blooms in the middle Vaal river: A site specific approach. Water Research 31(2), 271-279. 10.1016/s0043-1354(96)00248-5.

Cloot, A.H.J. and Piererse, A.J.H. 1999. Modelling phytoplankton in the Vaal river (South Africa). Water Science and Technology 40(10), 119-124. 10.1016/s0273-1223(99)00680-0.

Crossman, J., Bussi, G., Whitehead, P.G., Butterfield, D., Lannergård, E. and Futter, M.N. 2021. A New, Catchment-Scale Integrated Water Quality Model of Phosphorus, Dissolved Oxygen, Biochemical Oxygen Demand and Phytoplankton: INCA-Phosphorus Ecology (PEco). Water 13(5). 10.3390/w13050723.

de Souza Beghelli, F.G., Frascareli, D., Pompêo, M.L.M. and Moschini-Carlos, V. 2016. Trophic State Evolution over 15 Years in a Tropical Reservoir with Low Nitrogen Concentrations and Cyanobacteria Predominance. Water, Air, & Soil Pollution 227(3). 10.1007/s11270-016-2795-1.

Descy, J.-P. and Gosselain, V. 1994. Development and ecological importance of phytoplankton in a large lowland river (River Meuse, Belgium). Hydrobiologia 289(1-3), 139-155. 10.1007/bf00007415.
Descy, J.P., Leitao, M., Everbecq, E., Smitz, J.S. and Deliege, J.F. 2011. Phytoplankton of the River Loire, France: a biodiversity and modelling study. Journal of Plankton Research 34(2), 120-135. 10.1093/plankt/fbr085.
Descy, J.P., Servais, P., Smitz, J.S., Billen, G. and Everbecq, E. 1987. Phytoplankton biomass and production in the river meuse (Belgium). Water Research 21(12), 1557-1566. 10.1016/0043-1354(87)90141-2.
Dolgonosov, B.M., Korchagin, K.A. and Messineva, E.M. 2010. Statistical distributions of phytoplankton concentration in river water. Water Resources 37(2), 213-219. 10.1134/s0097807810020090.
Ducharne, A. 2008. Importance of stream temperature to climate change impact on water quality. Hydrology and Earth System Sciences 12(3), 797-810. 10.5194/hess-12-797-2008.
Dzialowski, A.R., Smith, V.H., Wang, S.-H., Martin, M.C. and Jr, F.d. 2011. Effects of non-algal turbidity on cyanobacterial biomass in seven turbid Kansas reservoirs. Lake and Reservoir Management 27(1), 6-14. 10.1080/07438141.2011.551027.
Even, S., Poulin, M., Mouchel, J.-M., Seidl, M. and Servais, P. 2004. Modelling oxygen deficits in the Seine River downstream of combined sewer overflows. Ecological Modelling 173(2-3), 177-196. 10.1016/j.ecolmodel.2003.08.019.
Everbecq, E., Gosselain, V., Viroux, L. and Descy, J.P. 2001. Potamon: a dynamic model for predicting phytoplankton composition and biomass in lowland rivers. Water Research 35(4), 901-912. 10.1016/s0043-1354(00)00360-2.
Gabyshev, V.A. 2021. Assessment of Potential Phytoplankton Transformations of Large Rivers of Eastern Siberia in Response to Global Climate Change. Biology Bulletin 48(6), 793-799. 10.1134/s106235902106008x.
Gamier, J., Billen, G. and Coste, M. 1995. Seasonal succession of diatoms and Chlorophyceae in the drainage network of the Seine River: Observation and modeling. Limnology and Oceanography 40(4), 750-765. 10.4319/lo.1995.40.4.0750.
Gao, Q., He, G., Fang, H., Bai, S. and Huang, L. 2018. Numerical simulation of water age and its potential effects on the water quality in Xiangxi Bay of Three Gorges Reservoir. Journal of Hydrology 566, 484-499. 10.1016/j.jhydrol.2018.09.033.
Ghermandi, A., Vandenberghe, V., Benedetti, L., Bauwens, W. and Vanrolleghem, P.A. 2009. Model-based assessment of shading effect by riparian vegetation on river water quality. Ecological Engineering 35(1), 92-104. 10.1016/j.ecoleng.2008.09.014.
Glaser, D., Rhea, J.R., Opdyke, D.R., Russell, K.T., Ziegler, C.K., Ku, W., Zheng, L. and Mastriano, J. 2009. Model of zebra mussel growth and water quality impacts in the Seneca River, New York. Lake and Reservoir Management 25(1), 49-72. 10.1080/07438140802714411.
Grover, J.P., Crane, K.W., Baker, J.W., Brooks, B.W. and Roelke, D.L. 2011. Spatial variation of harmful algae and their toxins in flowing-water habitats: a theoretical exploration. Journal of Plankton Research 33(2), 211-227. 10.1093/plankt/fbq070.

Grover, J.P., Hsu, S.B. and Wang, F.B.  2009.  Competition and coexistence in flowing habitats with a hydraulic storage zone. Mathematical Biosciences 222(1), 42-52. 10.1016/j.mbs.2009.08.006.
Grover, J.P., Roelke, D.L. and Brooks, B.W.  2017.  Population persistence in flowing-water habitats: Conditions where flow-based management of harmful algal blooms works, and where it does not. Ecological Engineering 99, 172-181. 10.1016/j.ecoleng.2016.11.044.
Guven, B. and Howard, A.  2006.  Modelling the growth and movement of cyanobacteria in river systems. Science of The Total Environment 368(2-3), 898-908. 10.1016/j.scitotenv.2006.03.035.
Guven, B. and Howard, A.  2007.  Identifying the critical parameters of a cyanobacterial growth and movement model by using generalised sensitivity analysis. Ecological Modelling 207(1), 11-21. 10.1016/j.ecolmodel.2007.03.024.
Guven, B. and Howard, A.  2011.  Sensitivity Analysis of a Cyanobacterial Growth and Movement Model under Two Different Flow Regimes. Environmental Modeling & Assessment 16(6), 577-589. 10.1007/s10666-011-9266-2.
Hajda, P. and Novotny, V.  1996.  Modelling impact of urban and upstream nonpoint sources on eutrophicationof the Milwaukee River. Water Science and Technology 33(4-5). 10.1016/0273-1223(96)00225-9.
Harvey, J.W., Choi, J. and Quion, K.  2024.  Metabolism Regimes in Regulated Rivers of the Illinois River Basin, USA. Scientific Data 11(1), 211. 10.1038/s41597-024-03037-1.
He, X., Wang, H., Fan, L., Liang, D., Ao, Y. and Zhuang, W.  2020.  Quantifying physical transport and local proliferation of phytoplankton downstream of an eutrophicated lake. Journal of Hydrology 585. 10.1016/j.jhydrol.2020.124796.
He, X., Wang, H., Yan, H. and Ao, Y.  2021a.  Numerical simulation of microcystin distribution in Liangxi River, downstream of Taihu Lake. Water Environment Research 93(10), 1934-1943. 10.1002/wer.1484.
He, X., Wang, H., Zhuang, W., Liang, D. and Ao, Y.  2021b.  Risk prediction of microcystins based on water quality surrogates: A case study in a eutrophicated urban river network. Environmental Pollution 275, 116651. 10.1016/j.envpol.2021.116651.
Heiskary, S.A. and Walker, W.W.  1995.  Establishing a ChlorophyllaGoal for a Run-of-the-river Reservoir. Lake and Reservoir Management 11(1), 67-76. 10.1080/07438149509354199.
Hesse, C. and Krysanova, V.  2016.  Modeling Climate and Management Change Impacts on Water Quality and In-Stream Processes in the Elbe River Basin. Water 8(2). 10.3390/w8020040.
Hong, D.-G., Kim, D.-K., Kim, H.-W., Do, Y., Lee, H.Y. and Joo, G.-J.  2016. Limnological assessment of the meteo-hydrological and physicochemical factors for summer cyanobacterial blooms in a regulated river system. Annales de Limnologie - International Journal of Limnology 52, 123-136. 10.1051/limn/2015038.
Honti, M., Istvanovics, V. and Kovacs, A.S.  2010.  Balancing between retention and flushing in river networks--optimizing nutrient management to improve trophic

state. Science of The Total Environment 408(20), 4712-4721. 10.1016/j.scitotenv.2010.06.054.

Hosseini, N., Chun, K. and Lindenschmidt, K.-E. 2016. Quantifying Spatial Changes in the Structure of Water Quality Constituents in a Large Prairie River within Two Frameworks of a Water Quality Model. Water 8(4). 10.3390/w8040158.

Hou, C., Chu, M.L. and Guzman, J.A. 2022. Risk assessment of harmful algal blooms (HAB) occurrence in the agroecosystem: A hydro-ecologic modeling framework and environmental risk matrix. Ecological Indicators 145(109617). 10.1016/j.ecolind.2022.109617.

Hou, G., Li, H., Recknagel, F. and Song, L. 2006. Modeling Phytoplankton Dynamics in the River Darling (Australia) Using the Radial Basis Function Neural Network. Journal of Freshwater Ecology 21(4), 639-647. 10.1080/02705060.2006.9664125.

Hou, G.X., Li, H.B., Recknagel, F. and Song, L.R. 2007. Using the moving window incorporated neural network to forecast the population behavior of *Nostocales* spp. in the River Darling, Australia. Fresenius Environmental Bulletin 16(3), 304-309.

Howard, A., Kirkby, M.J., Kneale, P.E. and McDonald, A.T. 2006. Modelling the growth of cyanobacteria (GrowSCUM). Hydrological Processes 9(7), 809-820. 10.1002/hyp.3360090707.

Hsu, S.-B., Wang, F.-B. and Zhao, X.-Q. 2013. Global dynamics of zooplankton and harmful algae in flowing habitats. Journal of Differential Equations 255(3), 265-297. 10.1016/j.jde.2013.04.006.

Hutchins, M.G., Johnson, A.C., Deflandre-Vlandas, A., Comber, S., Posen, P. and Boorman, D. 2010. Which offers more scope to suppress river phytoplankton blooms: reducing nutrient pollution or riparian shading? Science of The Total Environment 408(21), 5065-5077. 10.1016/j.scitotenv.2010.07.033.

Isabwe, A., Yang, J.R., Wang, Y., Liu, L., Chen, H. and Yang, J. 2018. Community assembly processes underlying phytoplankton and bacterioplankton across a hydrologic change in a human-impacted river. Science of The Total Environment 630, 658-667. 10.1016/j.scitotenv.2018.02.210.

Istvànovic, V. and Honti, M. 2011. Phytoplankton growth in three rivers: The role of meroplankton and the benthic retention hypothesis. Limnology and Oceanography 56(4), 1439-1452. 10.4319/lo.2011.56.4.1439.

Istvánovics, V., Honti, M., Kovács, Á., Kocsis, G. and Stier, I. 2014. Phytoplankton growth in relation to network topology: time-averaged catchment-scale modelling in a large lowland river. Freshwater Biology 59(9), 1856-1871. 10.1111/fwb.12388.

Jäger, C.G. and Borchardt, D. 2018. Longitudinal patterns and response lengths of algae in riverine ecosystems: A model analysis emphasising benthic-pelagic interactions. Journal of Theoretical Biology 442, 66-78. 10.1016/j.jtbi.2018.01.009.

Jeong, K.-S., Kim, D.-K., Jung, J.-M., Kim, M.-C. and Joo, G.-J. 2008. Non-linear autoregressive modelling by Temporal Recurrent Neural Networks for the prediction of freshwater phytoplankton dynamics. Ecological Modelling 211(3-4), 292-300. 10.1016/j.ecolmodel.2007.09.029.

Jeong, K.-S., Kim, D.-K., Whigham, P. and Joo, G.-J. 2003. Modelling *Microcystis aeruginosa* bloom dynamics in the Nakdong River by means of evolutionary computation and statistical approach. Ecological Modelling 161(1-2), 67-78. 10.1016/s0304-3800(02)00280-6.

Jia, H., Zhang, Y. and Guo, Y. 2010. The development of a multi-species algal ecodynamic model for urban surface water systems and its application. Ecological Modelling 221(15), 1831-1838. 10.1016/j.ecolmodel.2010.04.009.

Jung, W.S., Jo, B.G. and Kim, Y.D. 2023. A Study on the Occurrence Characteristics of Harmful Blue-Green Algae in Stagnant Rivers Using Machine Learning. Applied Sciences 13(6). 10.3390/app13063699.

Kim, D.-K., Cao, H., Jeong, K.-S., Recknagel, F. and Joo, G.-J. 2007. Predictive function and rules for population dynamics of Microcystis aeruginosa in the regulated Nakdong River (South Korea), discovered by evolutionary algorithms. Ecological Modelling 203(1-2), 147-156. 10.1016/j.ecolmodel.2006.03.040.

Kim, D., Jeong, K.-S., McKay, R.B., Chon, T.-S. and Joo, G.-J. 2012. Machine learning for predictive management: short and long term prediction of phytoplankton biomass using genetic algorithm based recurrent neural networks. International Journal of Environmental Research 6(1), 95-108. 10.22059/ijer.2011.476.

Kim, H.G., Hong, S., Jeong, K.-S., Kim, D.-K. and Joo, G.-J. 2019a. Determination of sensitive variables regardless of hydrological alteration in artificial neural network model of chlorophyll a: Case study of Nakdong River. Ecological Modelling 398, 67-76. 10.1016/j.ecolmodel.2019.02.003.

Kim, J., Jones, J.R. and Seo, D. 2021a. Factors affecting harmful algal bloom occurrence in a river with regulated hydrology. Journal of Hydrology: Regional Studies 33. 10.1016/j.ejrh.2020.100769.

Kim, J., Jung, W., An, J., Oh, H.J. and Park, J. 2023. Self-optimization of training dataset improves forecasting of cyanobacterial bloom by machine learning. Science of The Total Environment 866, 161398. 10.1016/j.scitotenv.2023.161398.

Kim, J., Kwak, J., Ahn, J.M., Kim, H., Jeon, J. and Kim, K. 2022a. Oscillation Flow Dam Operation Method for Algal Bloom Mitigation. Water 14(8). 10.3390/w14081315.

Kim, J., Lee, T. and Seo, D. 2017. Algal bloom prediction of the lower Han River, Korea using the EFDC hydrodynamic and water quality model. Ecological Modelling 366, 27-36. 10.1016/j.ecolmodel.2017.10.015.

Kim, J., Seo, D. and Jones, J.R. 2022b. Harmful algal bloom dynamics in a tidal river influenced by hydraulic control structures. Ecological Modelling 467. 10.1016/j.ecolmodel.2022.109931.

Kim, J.S., Seo, I.W. and Baek, D. 2018. Modeling spatial variability of harmful algal bloom in regulated rivers using a depth-averaged 2D numerical model. Journal of Hydro-environment Research 20, 63-76. 10.1016/j.jher.2018.04.008.

Kim, J.S., Seo, I.W. and Baek, D. 2019b. Seasonally varying effects of environmental factors on phytoplankton abundance in the regulated rivers. Scientific Reports 9(1), 9266. 10.1038/s41598-019-45621-1.

Kim, K.B., Uranchimeg, S. and Kwon, H.H. 2022c. A multivariate Chain-Bernoulli-based prediction model for cyanobacteria algal blooms at multiple stations in

South Korea. Environmental Pollution 313, 120078. 10.1016/j.envpol.2022.120078.

Kim, S., Kim, S., Mehrotra, R. and Sharma, A. 2020. Predicting cyanobacteria occurrence using climatological and environmental controls. Water Research 175, 115639. 10.1016/j.watres.2020.115639.

Kim, S., Mehrotra, R., Kim, S. and Sharma, A. 2021b. Assessing Countermeasure Effectiveness in Controlling Cyanobacterial Exceedance in Riverine Systems Using Probabilistic Forecasting Alternatives. Journal of Water Resources Planning and Management 147(10). 10.1061/(asce)wr.1943-5452.0001449.

Kim, S., Mehrotra, R., Kim, S. and Sharma, A. 2021c. Probabilistic forecasting of cyanobacterial concentration in riverine systems using environmental drivers. Journal of Hydrology 593. 10.1016/j.jhydrol.2020.125626.

Kim, T., Shin, J., Lee, D., Kim, Y., Na, E., Park, J.H., Lim, C. and Cha, Y. 2022d. Simultaneous feature engineering and interpretation: Forecasting harmful algal blooms using a deep learning approach. Water Research 215, 118289. 10.1016/j.watres.2022.118289.

Kireta, A.R., Reavie, E.D., Sgro, G.V., Angradi, T.R., Bolgrien, D.W., Hill, B.H. and Jicha, T.M. 2012. Planktonic and periphytic diatoms as indicators of stress on great rivers of the United States: Testing water quality and disturbance models. Ecological Indicators 13(1), 222-231. 10.1016/j.ecolind.2011.06.006.

Kobayashi, T., Ralph, T.J., Ryder, D.S. and Hunter, S.J. 2013. Gross primary productivity of phytoplankton and planktonic respiration in inland floodplain wetlands of southeast Australia: habitat-dependent patterns and regulating processes. Ecological Research 28(5), 833-843. 10.1007/s11284-013-1065-6.

Lee, D., Kim, M., Lee, B., Chae, S., Kwon, S. and Kang, S. 2022. Integrated explainable deep learning prediction of harmful algal blooms. Technological Forecasting and Social Change 185, 122046. 10.1016/j.techfore.2022.122046.

Lee, D.H., Fabian, P.S., Kim, J.H. and Kang, J.-H. 2021. HSPF-Based Assessment of Inland Nutrient Source Control Strategies to Reduce Algal Blooms in Streams in Response to Future Climate Changes. Sustainability 13(22). 10.3390/su132212413.

Lee, D.H., Kim, J.H., Park, M.-H., Stenstrom, M.K. and Kang, J.-H. 2020. Automatic calibration and improvements on an instream chlorophyll a simulation in the HSPF model. Ecological Modelling 415. 10.1016/j.ecolmodel.2019.108835.

Lee, S. and Lee, D. 2018. Four Major South Korea's Rivers Using Deep Learning Models. International Journal of Environmental Research 15(7). 10.3390/ijerph15071322.

Lévesque, D., Cattaneo, A., Hudon, C., Gagnon, P. and Weyhenmeyer, G. 2012. Predicting the risk of proliferation of the benthic cyanobacterium *Lyngbya wollei* in the St. Lawrence River. Canadian Journal of Fisheries and Aquatic Sciences 69(10), 1585-1595. 10.1139/f2012-087.

Lewis, D.M., Elliott, J.A., Lambert, M.F. and Reynolds, C.S. 2002. The simulation of an Australian reservoir using a phytoplankton community model: PROTECH. Ecological Modelling 150(1-2), 107-116. 10.1016/s0304-3800(01)00466-5.

Li, J., Li, D. and Wang, X. 2012. Three-dimensional unstructured-mesh eutrophication model and its application to the Xiangxi River, China. Journal of Environmental Sciences 24(9), 1569-1578. 10.1016/s1001-0742(11)60956-x.

Liao, A., Han, D., Song, X. and Yang, S. 2021. Impacts of storm events on chlorophyll-a variations and controlling factors for algal bloom in a river receiving reclaimed water. Journal of environmental management 297, 113376. 10.1016/j.jenvman.2021.113376.

Lindenschmidt, K.-E., Fleischbein, K. and Baborowski, M. 2007. Structural uncertainty in a river water quality modelling system. Ecological Modelling 204(3-4), 289-300. 10.1016/j.ecolmodel.2007.01.004.

Liu, W.C., Hsu, M.H. and Kuo, A.Y. 2000. Sensitivity analysis of a mathematical model of chlorophyll distribution in the tidal Keelung river. Journal of Environmental Science and Health, Part A 35(4), 489-514. 10.1080/10934520009376983.

Loewenthal, M., Neal, C., Whitehead, P.G., Wade, A.J. and Lázár, A.N. 2012. Reconciling observed and modelled phytoplankton dynamics in a major lowland UK river, the Thames. Hydrology Research 43(5), 576-588. 10.2166/nh.2012.029.

Loos, S., Shin, C.M., Sumihar, J., Kim, K., Cho, J. and Weerts, A.H. 2020. Ensemble data assimilation methods for improving river water quality forecasting accuracy. Water Research 171, 115343. 10.1016/j.watres.2019.115343.

Lucas, L.V., Thompson, J.K. and Brown, L.R. 2009. Why are diverse relationships observed between phytoplankton biomass and transport time? Limnology and Oceanography 54(1), 381-390. 10.4319/lo.2009.54.1.0381.

Lung, W.-S. 1988. The Role of Estuarine Modeling in Nutrient Control. Water Science and Technology 20(6-7), 243-252. 10.2166/wst.1988.0209.

Lung, W.S. 1986. Assessing Phosphorus Control in the James River Basin. Journal of Environmental Engineering 112(1), 44-60. 10.1061/(asce)0733-9372(1986)112:1(44).

Lv, X., Zhang, J., Liang, P., Zhang, X., Yang, K. and Huang, X. 2020. Phytoplankton in an urban river replenished by reclaimed water: Features, influential factors and simulation. Ecological Indicators 112. 10.1016/j.ecolind.2020.106090.

Maier, H.R. and Dandy, G.C. 1997. Modelling cyanobacteria (blue-green algae) in the River Murray using artificial neural networks. Mathematics and Computers in Simulation 43(3-6), 377-386. 10.1016/s0378-4754(97)00022-0.

Maier, H.R., Dandy, G.C. and Burch, M.D. 1998. Use of artificial neural networks for modelling cyanobacteria *Anabaena* spp. in the River Murray, South Australia. Ecological Modelling 105(2-3), 257-272. 10.1016/s0304-3800(97)00161-0.

Minaudo, C., Curie, F., Jullian, Y., Gassama, N. and Moatar, F. 2018. QUAL-NET, a high temporal-resolution eutrophication model for large hydrographic networks. Biogeosciences 15(7), 2251-2269. 10.5194/bg-15-2251-2018.

Mitrovic, S.M., Chessman, B.C., Bowling, L.C. and Cooke, R.H. 2006. Modelling suppression of cyanobacterial blooms by flow management in a lowland river. River Research and Applications 22(1), 109-114. 10.1002/rra.875.

Moreno-Ostos, E., Elliott, J.A., Cruz-Pizarro, L., Escot, C., Basanta, A. and George, D.G. 2007. Using a numerical model (PROTECH) to examine the impact of water transfers on phytoplankton dynamics in a Mediterranean reservoir. Limnetica 26(1), 1-11. 10.23818/limn.26.01.

Neres-Lima, V., Machado-Silva, F., Baptista, D.F., Oliveira, R.B.S., Andrade, P.M., Oliveira, A.F., Sasada-Sato, C.Y., Silva-Junior, E.F., Feijó-Lima, R., Angelini, R., Camargo, P.B. and Moulton, T.P. 2017. Allochthonous and autochthonous carbon flows in food webs of tropical forest streams. Freshwater Biology 62(6), 1012-1023. 10.1111/fwb.12921.

Nietch, C.T., Gains-Germain, L., Lazorchak, J., Keely, S.P., Youngstrom, G., Urichich, E.M., Astifan, B., DaSilva, A. and Mayfield, H. 2022. Development of a Risk Characterization Tool for Harmful Cyanobacteria Blooms on the Ohio River. Water 14(4), 1-23. 10.3390/w14040644.

Park, Y., Lee, H.K., Shin, J.K., Chon, K., Kim, S., Cho, K.H., Kim, J.H. and Baek, S.S. 2021. A machine learning approach for early warning of cyanobacterial bloom outbreaks in a freshwater reservoir. Journal of environmental management 288, 112415. 10.1016/j.jenvman.2021.112415.

Pathak, D., Hutchins, M., Brown, L., Loewenthal, M., Scarlett, P., Armstrong, L., Nicholls, D., Bowes, M. and Edwards, F. 2021. Hourly Prediction of Phytoplankton Biomass and Its Environmental Controls in Lowland Rivers. Water Resources Research 57(3). 10.1029/2020wr028773.

Pinckney, J.L., Millie, D.F., Vinyard, B.T. and Paerl, H.W. 1997. Environmental controls of phytoplankton bloom dynamics in the Neuse River Estuary, North Carolina, U.S.A. Canadian Journal of Fisheries and Aquatic Sciences 54(11), 2491-2501. 10.1139/f97-165.

Pyo, J., Cho, K.H., Kim, K., Baek, S.S., Nam, G. and Park, S. 2021. Cyanobacteria cell prediction using interpretable deep learning model with observed, numerical, and sensing data assemblage. Water Research 203, 117483. 10.1016/j.watres.2021.117483.

Pyo, J., Pachepsky, Y.A., Kim, M., Baek, S.-S., Lee, H., Cha, Y., Park, Y. and Cho, K.H. 2019. Simulating seasonal variability of phytoplankton in stream water using the modified SWAT model. Environmental Modelling & Software 122, 104073. 10.1016/j.envsoft.2017.11.005.

Pyo, J., Park, L.J., Pachepsky, Y., Baek, S.S., Kim, K. and Cho, K.H. 2020. Using convolutional neural network for predicting cyanobacteria concentrations in river water. Water Research 186, 116349. 10.1016/j.watres.2020.116349.

Qin, Q. and Shen, J. 2019. Physical transport processes affect the origins of harmful algal blooms in estuaries. Harmful Algae 84, 210-221. 10.1016/j.hal.2019.04.002.

Raimonet, M., Thieu, V., Silvestre, M., Oudin, L., Rabouille, C., Vautard, R. and Garnier, J. 2018. Landward Perspective of Coastal Eutrophication Potential Under Future Climate Change: The Seine River Case (France). Frontiers in Marine Science 5. 10.3389/fmars.2018.00136.

Rankinen, K., Cano Bernal, J.E., Holmberg, M., Vuorio, K. and Granlund, K. 2019. Identifying multiple stressors that influence eutrophication in a Finnish agricultural river. Science of The Total Environment 658, 1278-1292. 10.1016/j.scitotenv.2018.12.294.

Recknagel, F., French, M., Harkonen, P. and Yabunaka, K.-I. 1997. Artificial neural network approach for modelling and prediction of algal blooms. Ecological Modelling 96(1-3), 11-28. 10.1016/s0304-3800(96)00049-x.

Recknagel, F., Kim, D.K., Joo, G.J. and Cao, H. 2017. Response of Microcystis and Stephanodiscus to Alternative Flow Regimes of the Regulated River Nakdong (South Korea) Quantified By Model Ensembles Based on the Hybrid Evolutionary Algorithm (HEA). River Research and Applications 33(6), 949-958. 10.1002/rra.3141.

Remmal, Y., Hudon, C., Hamilton, P.B., Rondeau, M. and Gagnon, P. 2017. Forecasting the magnitude and composition of phytoplankton blooms in a eutrophic lowland river (Rivière Yamaska, Que., Canada). Canadian Journal of Fisheries and Aquatic Sciences 74(8), 1298-1311. 10.1139/cjfas-2016-0305.

Rode, M., Poser, K. and Lindenschmidt, K.E. 2005. Impact of morphological parameters on water quality variables of a regulated lowland river. Water Science and Technology 52(6), 187-193. 10.2166/wst.2005.0167.

Rose, A.K., Kinder, J.E., Fabbro, L. and Kinnear, S. 2019. A phytoplankton risk matrix: combining health, treatment, and aesthetic considerations in drinking water supplies. Environment Systems and Decisions 39(2), 163-182. 10.1007/s10669-018-9711-8.

Rounds, S.A. and Wood, T.M. 2001. Modeling water quality in the Tualatin River, Oregon, 1991-1997. U.S. Geological Survey Water Resources Investigations Report 01-4041, 53 p. 10.3133/wri014041.

Rounds, S.A., Wood, T.M. and Lynch, D.D. 1999. Modeling discharge, temperature, and water quality in the Tualatin River, Oregon. U.S. Geological Survey Water-Supply Paper 2465-B. 10.3133/wsp2465B.

Savoy, P. and Harvey, J.W. 2023. Predicting Daily River Chlorophyll Concentrations at a Continental Scale. Water Resources Research 59(11), e2022WR034215. 10.1029/2022wr034215.

Scharfe, M., Callies, U., Blöcker, G., Petersen, W. and Schroeder, F. 2009. A simple Lagrangian model to simulate temporal variability of algae in the Elbe River. Ecological Modelling 220(18), 2173-2186. 10.1016/j.ecolmodel.2009.04.048.

Schöl, A., Hein, B., Wyrwa, J. and Kirchesch, V. 2014. Modelling water quality in the Elbe and its estuary–Large scale and long term applications with focus on the oxygen budget of the estuary. Die Küste 81(495), 203-232.

Schöl, A., Kirchesch, V., Bergfeld, T., Schöll, F., Borcherding, J. and Müller, D. 2002. Modelling the Chlorophyll a Content of the River Rhine -Interrelation between Riverine Algal Production and Population Biomass of Grazers, Rotifers and the Zebra Mussel, *Dreissena polymorpha*. International Review of Hydrobiology 87(2-3), 295-317. 10.1002/1522-2632(200205)87:2/3<295::Aid-iroh295>3.0.Co;2-b.

Seo, D.-I., Kim, M.-A. and Ahn, J.-H. 2012. Prediction of Chlorophyll-a Changes due to Weir Constructions in the Nakdong River Using EFDC-WASP Modelling. Environmental Engineering Research 17(2), 95-102. 10.4491/eer.2012.17.2.095.

Seo, D. and Song, Y. 2015. Application of three-dimensional hydrodynamics and water quality model of the Youngsan River, Korea. Desalination and Water Treatment 54(13), 3712-3720. 10.1080/19443994.2014.923210.

Shan, K., Ouyang, T., Wang, X., Yang, H., Zhou, B., Wu, Z. and Shang, M. 2022. Temporal prediction of algal parameters in Three Gorges Reservoir based on highly time-resolved monitoring and long short-term memory network. Journal of Hydrology 605, 127304. 10.1016/j.jhydrol.2021.127304.

Shin, J., Yoon, S. and Cha, Y. 2017. Prediction of cyanobacteria blooms in the lower Han River (South Korea) using ensemble learning algorithms. Desalination and Water Treatment 84, 31-39. 10.5004/dwt.2017.20986.

Shin, J., Yoon, S., Kim, Y., Kim, T., Go, B. and Cha, Y. 2021. Effects of class imbalance on resampling and ensemble learning for improved prediction of cyanobacteria blooms. Ecological Informatics 61. 10.1016/j.ecoinf.2020.101202.

Sipkay, C., Kiss-Keve, T., Vadadi-Fülöp, C., Homoródi, R. and Hufnagel, L. 2012. Simulation modeling of phytoplankton dynamics in a large eutrophic river, Hungary — Danubian Phytoplankton Growth Model (DPGM). Biologia 67(2), 323-337. 10.2478/s11756-012-0004-2.

Smith, E.A., Akasapu-Smith, M., Petkewich, M.D. and Conrads, P.A. 2022. Evaluation of the Bushy Park Reservoir Three-Dimensional Hydrodynamic and Water-Quality Model, South Carolina, 2012–15. U.S. Geological Survey Open-File Report 2022-1079. https://doi.org/10.3133/ofr20221079.

Son, G., Kim, D., Kim, Y.D., Lyu, S. and Kim, S. 2020. A Forecasting Method for Harmful Algal Bloom (HAB)-Prone Regions Allowing Preemptive Countermeasures Based only on Acoustic Doppler Current Profiler Measurements in a Large River. Water 12(12), 3488. 10.3390/w12123488.

Song, Y., Chen, M., Li, J., Zhang, L., Deng, Y. and Chen, J. 2023. Can selective withdrawal control algal blooms in reservoirs? The underlying hydrodynamic mechanism. Journal of Cleaner Production 394. 10.1016/j.jclepro.2023.136358.

Stringfellow, W., Herr, J., Litton, G., Brunell, M., Borglin, S., Hanlon, J., Chen, C., Graham, J., Burks, R., Dahlgren, R., Kendall, C., Brown, R. and Quinn, N. 2009. Investigation of river eutrophication as part of a low dissolved oxygen total maximum daily load implementation. Water Science and Technology 59(1), 9-14. 10.2166/wst.2009.739.

Su, Y., Hu, M., Wang, Y., Zhang, H., He, C., Wang, Y., Wang, D., Wu, X., Zhuang, Y., Hong, S. and Trolle, D. 2022. Identifying key drivers of harmful algal blooms in a tributary of the Three Gorges Reservoir between different seasons: Causality based on data-driven methods. Environmental Pollution 297, 118759. 10.1016/j.envpol.2021.118759.

Sullivan, A.B. and Rounds, S.A. 2018. Modeling hydrodynamics, water temperature, and water quality in Klamath Straits Drain, Oregon and California, 2012–15. U.S. Geological Survey Scientific Investigations Report. 10.3133/sir20185134.

Sullivan, A.B., Rounds, S.A., Deas, M.L., Asbill, J.R., Wellman, R.E., Stewart, M.A., Johnston, M.W. and Sogutlugil, I.E. 2011. Modeling hydrodynamics, water temperature, and water quality in the Klamath River upstream of Keno Dam, Oregon, 2006-09. U.S. Geological Survey Scientific Investigations Report. 10.3133/sir20115105.

Teles, L.O., Vasconcelos, V., Pereira, E. and Saker, M. 2006. Time series forecasting of cyanobacteria blooms in the Crestuma Reservoir (Douro River, Portugal) using artificial neural networks. Environmental Management 38(2), 227-237. 10.1007/s00267-005-0074-9.

Thebault, J.-M. and Qotbi, A. 1999. A model of phytoplankton development in the Lot River (France). Water Research 33(4), 1065-1079. 10.1016/s0043-1354(98)00284-x.

Vis, C., Hudon, C., Carignan, R. and Gagnon, P. 2007. Spatial Analysis of Production by Macrophytes, Phytoplankton and Epiphyton in a Large River System under Different Water-Level Conditions. Ecosystems 10(2), 293-310. 10.1007/s10021-007-9021-3.

Wan, L., Wang, X.H. and Peirson, W. 2022. Impacts of Climate Change and Non-Point-Source Pollution on Water Quality and Algal Blooms in the Shoalhaven River Estuary, NSW, Australia. Water 14(12). 10.3390/w14121914.

Wang, B., Qiu, X.-L., Peng, X. and Wang, F. 2018a. Phytoplankton community structure and succession in karst cascade reservoirs, SW China. Inland Waters 8(2), 229-238. 10.1080/20442041.2018.1443550.

Wang, F.-B. 2015. A PDE system modeling the competition and inhibition of harmful algae with seasonal variations. Nonlinear Analysis: Real World Applications 25, 258-275. 10.1016/j.nonrwa.2015.02.010.

Wang, F.-B., Hsu, S.-B. and Zhao, X.-Q. 2015. A reaction–diffusion–advection model of harmful algae growth with toxin degradation. Journal of Differential Equations 259(7), 3178-3201. 10.1016/j.jde.2015.04.018.

Wang, J. and Zhang, Z. 2020. Phytoplankton, dissolved oxygen and nutrient patterns along a eutrophic river-estuary continuum: Observation and modeling. Journal of environmental management 261, 110233. 10.1016/j.jenvman.2020.110233.

Wang, L., Xie, Y., Xu, J., Zhang, H., Wang, X., Yu, J., Sun, Q., Zhao, Z., Elhoseny, M. and Yuan, X. 2019a. Prediction method of cyanobacterial blooms spatial-temporal sequence based on deep belief network and fuzzy expert system. Journal of Intelligent & Fuzzy Systems 38(2), 1487-1498. 10.3233/jifs-179512.

Wang, S., Flipo, N. and Romary, T. 2018b. Time-dependent global sensitivity analysis of the C-RIVE biogeochemical model in contrasted hydrological and trophic contexts. Water Research 144, 341-355. 10.1016/j.watres.2018.07.033.

Wang, S., Flipo, N. and Romary, T. 2019b. Oxygen data assimilation for estimating micro-organism communities' parameters in river systems. Water Research 165, 115021. 10.1016/j.watres.2019.115021.

Wang, Z., Chai, F., Dugdale, R., Liu, Q., Xue, H., Wilkerson, F., Chao, Y., Zhang, Y. and Zhang, H. 2020. The interannual variabilities of chlorophyll and nutrients in San Francisco Bay: a modeling study. Ocean Dynamics 70(8), 1169-1186. 10.1007/s10236-020-01386-0.

Wang, Z., Wang, H., Shen, J., Ye, F., Zhang, Y., Chai, F., Liu, Z. and Du, J. 2019c. An analytical phytoplankton model and its application in the tidal freshwater James River. Estuarine, Coastal and Shelf Science 224, 228-244. 10.1016/j.ecss.2019.04.051.

Waylett, A.J., Hutchins, M.G., Johnson, A.C., Bowes, M.J. and Loewenthal, M. 2013. Physico-chemical factors alone cannot simulate phytoplankton behaviour in a lowland river. Journal of Hydrology 497, 223-233. 10.1016/j.jhydrol.2013.05.027.

Welker, M. and Walz, N. 1999. Plankton dynamics in a river-lake system–on continuity and discontinuity. Hydrobiologia 408(0), 233-239. 10.1007/978-94-017-2986-4_25.
Whitehead, P.G., Bussi, G., Bowes, M.J., Read, D.S., Hutchins, M.G., Elliott, J.A. and Dadson, S.J. 2015. Dynamic modelling of multiple phytoplankton groups in rivers with an application to the Thames river system in the UK. Environmental Modelling & Software 74, 75-91. 10.1016/j.envsoft.2015.09.010.
Wu, T., Luo, L., Qin, B., Cui, G., Yu, Z. and Yao, Z. 2009. A vertically integrated eutrophication model and its application to a river-style reservoir--Fuchunjiang, China. Journal of Environmental Sciences 21(3), 319-327. 10.1016/s1001-0742(08)62271-8.
Xu, S., He, G., Fang, H., Bai, S. and Wu, X. 2022. Parameter uncertainty and sensitivity analysis of the three Gorges Reservoir and Xiangxi River EFDC model. Journal of Hydrology 610. 10.1016/j.jhydrol.2022.127881.
Yan, H., Wang, H., He, X., Liu, Y., Tang, Q., Yang, Y. and Yuan, W. 2021. Transverse distribution of cyanobacteria in a regulated urban river. Ecohydrology 14(3). 10.1002/eco.2274.
Yi, H.S., Park, S., An, K.G. and Kwak, K.C. 2018. Algal Bloom Prediction Using Extreme Learning Machine Models at Artificial Weirs in the Nakdong River, Korea. International Journal of Environmental Research 15(10). 10.3390/ijerph15102078.
Yoshioka, H. and Yaegashi, Y. 2017. Robust stochastic control modeling of dam discharge to suppress overgrowth of downstream harmful algae. Applied Stochastic Models in Business and Industry 34(3), 338-354. 10.1002/asmb.2301.
Zhao, X., Zhang, H. and Tao, X. 2013. Predicting the short-time-scale variability of chlorophyll a in the Elbe River using a Lagrangian-based multi-criterion analog model. Ecological Modelling 250, 279-286. 10.1016/j.ecolmodel.2012.11.018.
Ziemińska-Stolarska, A. and Kempa, M. 2021. Modeling and Monitoring of Hydrodynamics and Surface Water Quality in the Sulejów Dam Reservoir, Poland. Water 13(3). 10.3390/w13030296.
